# Molecular characterization of response to etrolizumab and anti-TNF reveals treatment resistance in ulcerative colitis is associated with an abundance of residual neutrophil subsets and inflammatory fibroblast populations

**DOI:** 10.1101/2024.07.02.601267

**Authors:** Shadi Toghi Eshghi, John Mark Gubatan, Parisa Mazrooei, Luis Quintanilla, Allen Nguyen, Amelia Au-Yeung, Derek Rudolph Holman, Chikara Takahashi, Courtney Schiffman, William O’Gorman, Mary Keir, Saroja Ramanujan, Stephan Rogalla, Jason A. Hackney, Jacqueline M McBride

## Abstract

Ulcerative colitis (UC) is a chronic inflammatory large bowel disease characterized by immune cell infiltration and continuous erosion of intestinal crypts, causing severe ulceration and abdominal pain. In the etrolizumab Phase 3 studies, transcriptional analyses of colonic biopsies revealed reductions in genes associated with aEb7+ intraepithelial lymphocytes with etrolizumab but not adalimumab. Both treatments significantly reduced stromal and myeloid cell-associated genes, with changes associated with MCS remission status. Generation of a single-cell atlas from inflamed and uninflamed colonic biopsies from UC patients led to the identification of thirty-six discrete cell populations, including cells of the myeloid compartment. The UC atlas was used to generate cell-specific signatures, allowing for cellular deconvolution of the Phase 3 datasets. It revealed significant reductions in neutrophil subsets, monocyte-derived macrophages, and inflammatory fibroblasts, as well as increases in colonic epithelial cells common to both etrolizumab and adalimumab. Pseudo-time trajectory analyses identified four unique neutrophil subsets with unique cell phenotypes reflecting changes in cell state or differentiation from PADI4hi, OSMhi, MX1hi, and ultimately to CXCR4hi populations. PADI4hi and OSMhi neutrophils exhibited high levels of proteases (MMP9, LYZ), inflammatory cytokines (CXCL1, IL1B, OSM), and abundant cytokine or chemokine receptors (CXCR1, CXCR2). MX1 populations expressed markers indicating prior IFN exposure (MX1, IFIT1).

In contrast, more differentiated or mature neutrophils exhibited high levels of CXCL2, TNF-a, and CXCR4, as well as angiogenic factors like VEGFA. PADI4hi and OSMhi neutrophils, we predict, have abundant cytokine and chemokine interactions with inflammatory fibroblasts within the inflamed colon, such as OSM: OSMR and IL1B: IL1R1 interactions. Changes in PADI4hi and OSMhi neutrophils were closely associated with MCS remission in both etrolizumab and adalimumab-treated patients. In contrast, only minor changes in CXCR4hi neutrophils were observed and not associated with clinical outcomes. Our results suggest that neutrophils are not only heterogeneous in phenotype but have abundant cell-cell interactions in inflamed colonic tissue that are likely implicated in maintaining chronic disease activity. We hypothesize that limiting the interactions between neutrophils and other myeloid cells with resident cells such as inflammatory fibroblasts may reduce the production of inflammatory mediators and limit activation and infiltration of neutrophils, which may be necessary for achieving greater rates of clinical remission in response to interventional agents.

## Introduction

Inflammatory bowel disease (IBD), including ulcerative colitis (UC), is a chronic inflammatory disorder of the gastrointestinal tract that has emerged as a globally relevant disease with increasing incidence worldwide and associated with significant morbidity and healthcare utilization (1–3). Although biologic therapies with diverse mechanisms of action (e.g., anti-TNF, anti-integrin, anti-cytokines, JAK inhibitors) have expanded the therapeutic armamentarium of IBD, only about 30-40% of patients respond to biologics and a significant proportion lose response over time (4, 5). There remains an unmet clinical need to understand the mechanisms of biologic therapy failure in IBD. Identifying and targeting the specific cell types, cellular interactions, and cellular pathways that mediate response or resistance to biologics could lead to more effective and precise therapeutic strategies to improve clinical outcomes in patients with IBD.

Etrolizumab is a fully humanized monoclonal anti-integrin beta 7 (b7) antibody with a dual mechanism of action: blocking mucosal homing gut homing of a4b7-bearing lymphocytes and reducing retention of aEb7-bearing intraepithelial lymphocytes (6–8). Results from Ph3 studies of etrolizumab had mixed clinical outcomes (6–9). Among those, HIBISCUS I and II were two identical randomized, double-blind, placebo-controlled studies with head-to-head efficacy comparisons with adalimumab conducted in patients with moderately to severely active ulcerative colitis and naive to biologic treatment (9). In the HIBISCUS I study, Etrolizumab showed a higher proportion of patients achieving remission during induction treatment relative to placebo; however, this effect was not replicated in HIBISCUS II. In prior studies, evidence suggested an impact on the frequency of aEb7 cells and reductions in infiltrating CD8+ T cells (6,7). The complete effect of etrolizumab in UC patients has not been described, and given the mixed Ph3 clinical outcomes, it was of interest to understand changes in mucosal expression profiles of responder and non-responder patients with direct comparisons to an effective treatment such as adalimumab.

Several reports describe transcriptional programs associated with disease severity, response to treatment, and non-response to biologics (10–17). Both immune and non-immune cells have been implicated, and through the advancement of single-cell and spatial methods, the evidence is mounting, pointing to the important role of myeloid cells in disease and response (13,18). It is widely appreciated in IBD that the presence and, importantly, the persistence of neutrophils is a critical feature linked to mucosal inflammation as well as a vital component found in Geboes, Nancy, and Robart’s histological indices used to grade histological disease severity or deep remission, however, only recently datasets capturing neutrophils emerged (18). Furthermore, neutrophil products such as fecal calprotectin have served as valuable biomarkers reflecting intestinal inflammation, and guidance suggests, combinations of symptoms and biomarkers, could be used as a non-invasive tool to monitor endoscopic disease in patients (19).

We sought to characterize immune and non-immune populations, including inflammatory macrophages and neutrophils, and their interactions with resident cells to better characterize the relationship of changes in these populations within the etrolizumab Ph3 studies. The work herein comprehensively evaluates the transcriptional profiles of immune cells with single-cell granularity and characterizes their phenotype in moderate-severe UC patients with comparisons between matched uninflamed and inflamed tissue. Importantly, we assessed transcriptional changes within our etrolizumab and adalimumab-treated UC patients within the HIBISCUS studies by leveraging UC single-cell datasets to deconvolve the on-treatment changes, especially within the myeloid compartment to assess cellular changes on treatment and relationships to response outcomes. Importantly, this study reports the most in-depth characterization of neutrophils in UC at the single-cell level and, through the application to clinical outcomes, provides a better understanding of the roles of neutrophils in response to treatment.

## Results

### Etrolizumab treatment significantly changes integrin gene expression and genes associated with inflammatory and epithelial cells

Given the mechanism of action of etrolizumab, a targeted assessment of integrin gene expression was performed to determine any relative changes in the homing of b7-expressing cells, including a4b7+ and aEb7+ cell populations. Baseline expression values for each of the integrins were similar across all three treatment groups, and a significant reduction was observed for ITGA4 and ITGB7 but not ITGB1 after both etrolizumab and adalimumab treatment (Figure 1A-D and Figure S1A-D). In contrast, ITGAE expression was reduced only with etrolizumab. No significant changes were observed with placebo treatment for these integrin genes (Figure S1 E-H). Unbiased transcriptome-wide analysis revealed that >1800 genes were significantly changed relative to baseline after etrolizumab (1261 genes decreased, 347 genes increased) or adalimumab treatment (1137 genes decreased, 112 genes increased) with changes of at least 1.5 fold relative to baseline and an FDR of 0.05. (Figure 1E-F, Table S1). The majority of genes reduced on treatment are associated with immune cell subsets and inflammatory processes (Table S2). Genes with increased expression were primarily expressed by or associated with functions of epithelial cell populations (Table S2).

**Figure 1.**
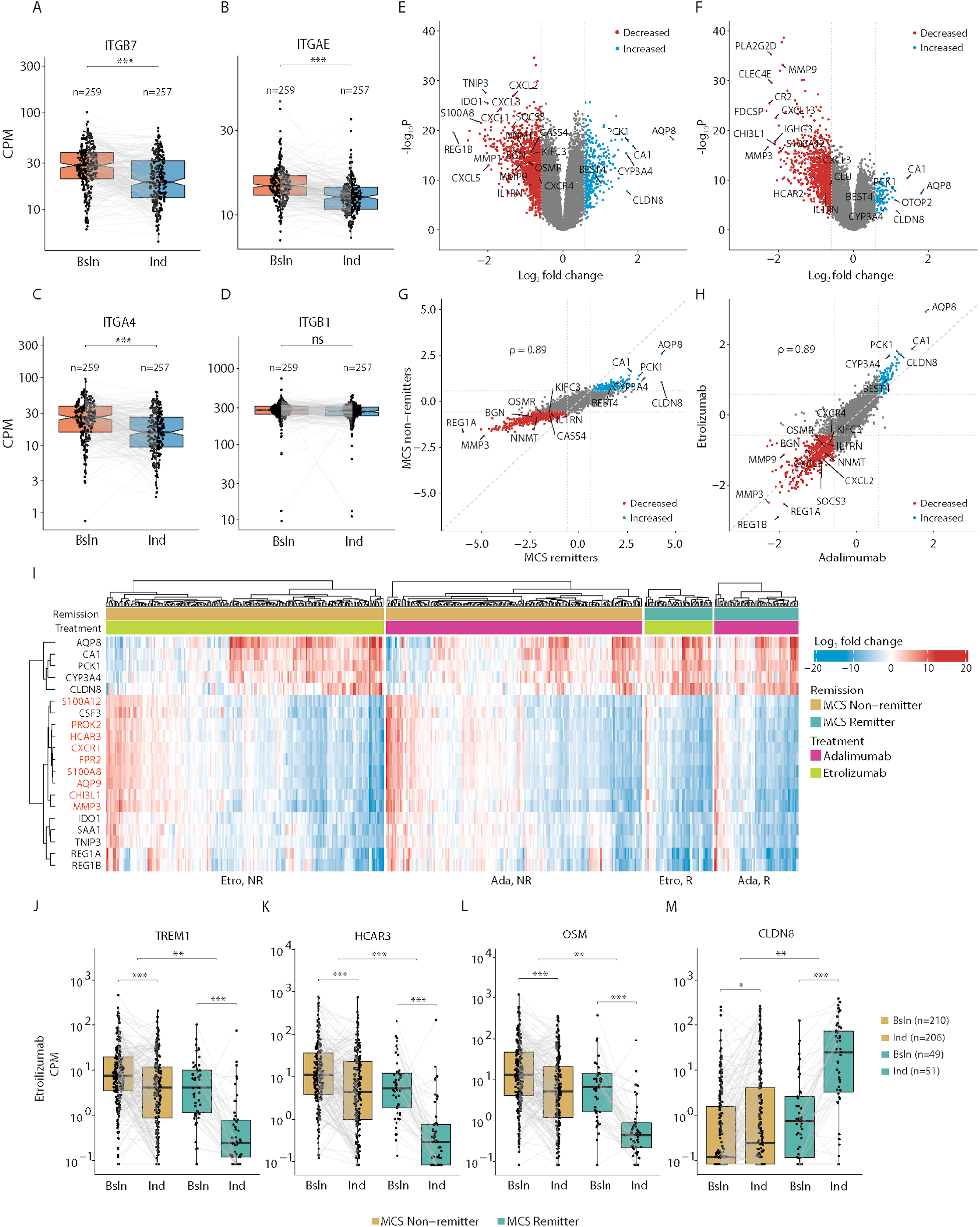
Transcriptomic analysis of etrolizumab, adalimumab, and placebo treatment in colonic biopsies. **A.-D.** Expression of selected integrins before (baseline) and after (induction; week 10) etrolizumab treatment. Each point represents an individual patient, and pre-and post-treatment trajectories for each patient are shown (Gray bar). Expression values are in normalized counts per million (CPM). Box plots represent the upper and lower quartiles, the middle line represents the median, notches show 1.58x the interquartile range (IQR) divided by the square root of the number of samples measured, and the whiskers extend to the most extreme point no more than 1.5x outside of the interquartile ranges. **E.-F.** Volcano plots showing log2 fold change on the x-axis and -log10 p-value on the y-axis for changes in gene expression after etrolizumab (**E**) or adalimumab (**F**) treatment for 10 weeks. Each point represents a gene, colored blue for genes significantly up-regulated after treatment (fold change > 1.5x at an FDR of 0.05) and colored red for genes significantly down-regulated after treatment (fold change < −1.5x at an FDR of 0.05). **G.** Comparison of log2 fold changes after etrolizumab treatment in either remitters (x-axis) or non-remitters (y-axis). Each point represents a gene, colored as in **E** above. Spearman correlation coefficient comparing the log2 fold changes across the conditions is shown. **H.** Comparison of log2 fold changes between adalimumab (x-axis) and etrolizumab (y-axis). Each point represents a gene, colored as in **E** above. Spearman correlation coefficient comparing the log2 fold changes across the conditions is shown. **I.** Heatmap of log2 fold changes of top differentially expressed genes after etrolizumab treatment. Each row shows a gene, and each column shows the difference in expression for a patient between week 10 and baseline measures. Patients are grouped by treatment arm and remission status. Labels from genes previously reported to be expressed by myeloid cells are shown in orange. **J.-M.** Expression of selected neutrophil-associated or epithelium-associated genes before (baseline) or after (induction; week 10) etrolizumab treatment. Each point represents a sample collected from a patient at the respective time point, and patient measurements are joined by a line. Boxes are colored by the remission status of the patient at week 10. Boxes represent the upper and lower quartile, the middle line represents the median, and the whiskers extend to the most extreme point no more than 1.5x the interquartile range from the box boundaries. For all panels, stars indicate Benjamini-Hochberg corrected p-values: ns: p > 0.01; *: p < 0.01; ** p < 0.001; *** p < 0.0001.

Changes in gene expression after etrolizumab treatment were directionally similar between MCS non-remitters and remitters; however, the magnitude of change was significantly greater in MCS remitters (Figure 1G). Likewise, gene expression changes were highly concordant between etrolizumab and adalimumab, with a strong correlation (Spearman rho = 0.89) between effect sizes for each treatment (Figure 1H and Table S1). Patients in the placebo arm showed directionally similar changes in gene expression, with reduced magnitude, and only a modest number of genes showed a statistically significant treatment effect (179 genes decreased and 43 genes increased at least 1.5 fold, with an FDR of 0.05; Figure S1I-K).

Upon closer examination of differentially expressed genes (DEGs) with the highest magnitude of change, the most striking enrichment of genes appeared to be associated with myeloid lineage cells, including neutrophils (Figure 1I). At an individual gene level, examples of the top twenty most differentially expressed genes included TREM1, HCAR3, and OSM, which were all significantly down-regulated after etrolizumab (Figure 1J-L) or adalimumab treatment (Figure S1L-N), with a substantially more significant change found in remitters in comparison to non-remitters. By contrast, epithelial genes such as CLDN8 were increased after etrolizumab or adalimumab treatment, with an increased magnitude observed in remitters compared to non-remitters (Figure 1M, S10). Though directionally consistent trends were observed, these genes were not significantly modulated in placebo-treated patients (Figure S1P-S).

### Identification of immune and non-immune cell populations in an expanded UC single-cell RNA-Seq Atlas

Given the high proportion of myeloid-derived genes being differentially expressed within bulk RNA sequencing from colonic tissue and the association of that lineage with disease activity, further analyses were performed to deeply characterize the sub-populations within each major cell types of interest and to understand changes in gene expression related to disease activity. We collected biopsies from inflamed and uninflamed colonic tissue from 20 moderate to severe UC patients, including 18 sets of matched pairs (Figure 2A, Table S3). In aggregate, we generated data from 408,930 cells, of which 236,855 were derived from inflamed colonic tissue and 172,075 cells from uninflamed tissue. Using a semi-automated annotation approach (see Methods for details), seventeen major cell populations were identified, encompassing stromal, epithelial, adaptive, and innate immune cells, including 47,516 neutrophils (Figure 2B, Figure S2A). Additionally, increased inflammatory cells, including neutrophils, were observed in biopsy samples collected from the inflamed area, and regions scored as three for endoscopic score (Figure S2B). Using more detailed annotation, we identified a total of thirty-six cell subsets including T cell subsets (CD4+ T helper, CD8+ cytotoxic, KI67+ cycling T cells), B cell subsets (IGHD+FCER2+ Follicular, MS4A1+AICDA+MKI67- germinal center, MKI67+PCNA+ cycling, MZB1+XBP1+PRDM1+ plasma cells/plasmablasts), myeloid subsets (VCAN+ monocyte-derived macrophages (MD macrophage), C1QA+ resident macrophages, CD1C+ dendritic cells, VNN2+ neutrophils, KIT+ mast cells, CLC+ eosinophils), IL7+ ILCs, NKG7+ NK cells, endothelial cells (ACKR1+, CD36+, CXCL12+ subsets), S100B+ glial cells, RGS5+ pericytes, HBB+ erythrocytes, epithelial cells (CA1+CA2+ enterocytes, BEST4+ enterocytes, CHGA+ enteroendocrine, MUC2+ goblet, SH2D6+ tuft cells, PCNA+LGR5+ stem cells, TOP2A+ transit amplifying), ACTA2+ SMC, and fibroblasts (PI6+ adventitial, WNT2B+ crypt-bottom, PDGFRA+ crypt-top, ADAMDEC1+ABCA8+ lamina propria, IL11+ inflammatory) (Figure 2C, S3A).

**Figure 2.**
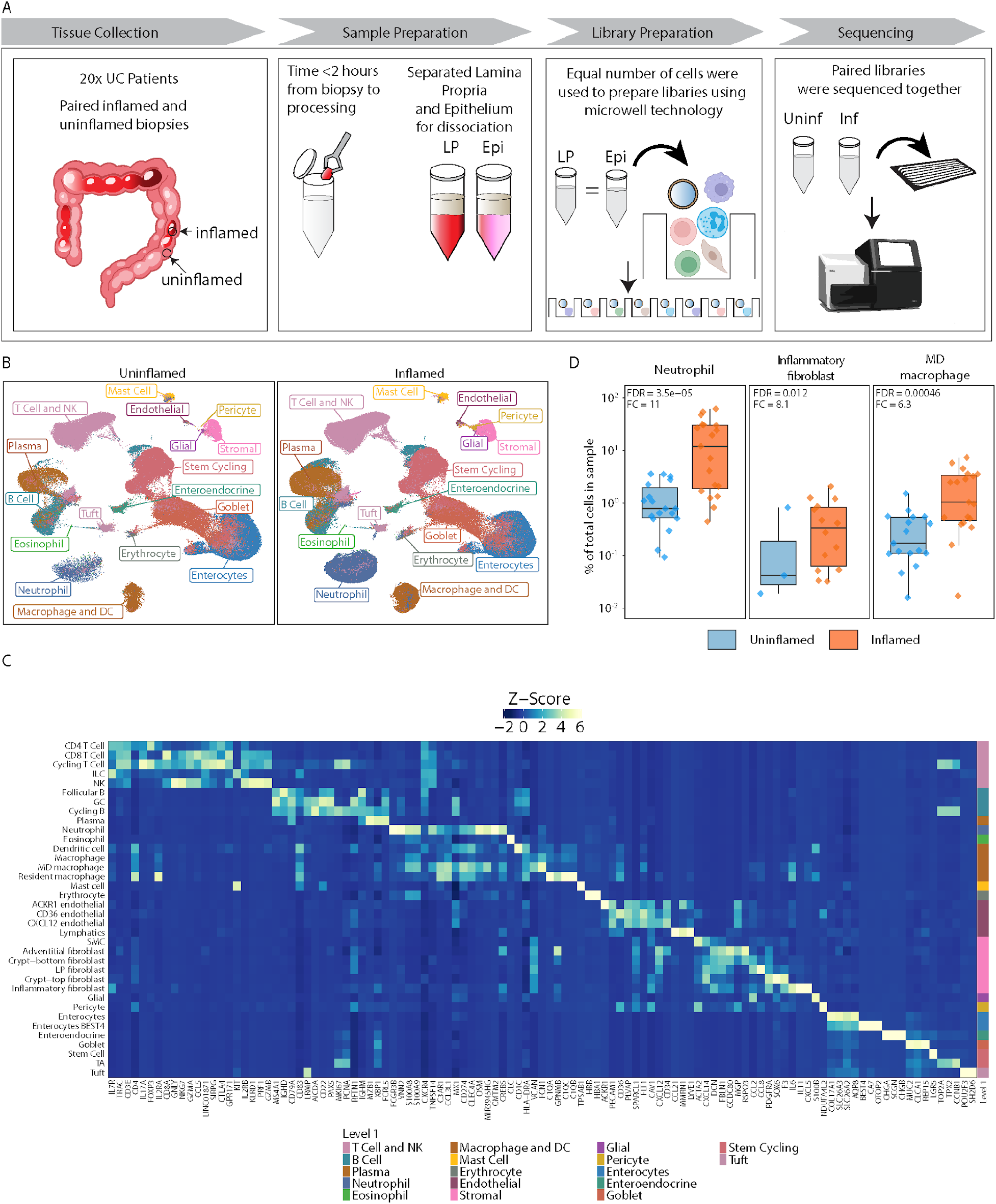
Generation of an ulcerative colitis single-cell atlas that captures tissue neutrophils. **A.** Experimental design. Biopsies were collected from both inflamed and uninflamed tissue from ulcerative colitis patients. To retain sensitive cells, including neutrophils, sample processing time was <2 hours from biopsy collection. We used the Rhapsody microwell platform to collect whole transcriptome data. **B.** UMAP of cells annotated at the broadest classification level. Each point represents an individual cell derived from uninflamed tissues (left panel) or inflamed colonic tissue (right panel), with each cell population colored by the lineage. **C.** Relative abundance of differentially abundant lineages. Relative abundance is shown as a percentage of total cells in each sample for neutrophils, inflammatory fibroblasts, and monocyte-derived macrophages across inflamed and uninflamed samples (Neutrophil, n=18 uninflamed, 19 inflamed) (Inflammatory Fibroblast, n = 3 uninflamed, 14 inflamed), (Monocyte-derived macrophage, n =17 uninflamed, 19 inflamed). Fold difference and FDR values were calculated as described in the methods. **D.** Heatmap of marker genes for intermediate-level annotations. Columns show individual genes specific to an individual cell type or broader lineage. Colors represent the centered and scaled pseudo-bulk expression levels. Broad lineage classification is demonstrated by the color bar on the right.

Importantly, this single-cell atlas allowed for the identification of genes associated with the myeloid lineages of interest, including macrophages and neutrophils, which have been consistently associated with resistance to biologic therapies (13, 20, 21) (Figure 1G). Assessment of individual cell subsets showed that, amongst all cell subsets identified, significant differences in relative abundance (negative binomial test, FDR < 0.05) were evident with elevations in neutrophils, MD macrophages, inflammatory fibroblasts, pericytes, ACKR1+ endothelial cells, and reductions in BEST4+ enterocytes and adventitial fibroblasts (Figure 2D, S2C, S2D).

### Cellular context of gene expression changes with disease and treatment

We used this single-cell atlas to generate cell-specific gene modules to deconvolve the clinical transcriptomic data and allow further elucidation of cellular changes found in etrolizumab- and adalimumab-treated patients. Each derived gene module was intended to be specific to a given cell population and sub-populations of a related cellular lineage (see Methods for details) while minimizing the overlap between related and unrelated populations (Figure S3A, Table S4). Examining bulk RNA-sequencing data derived from pooled HIBISCUS studies, it was evident there was a strong correlation between expression between most immune and stromal populations and a strong anti-correlation of their expression with epithelial cell subsets, regardless of treatment (Figure 3A, S4A). Higher expression markers of both immune cells, such as macrophages, neutrophils, and T cells, and stromal populations, such as pericytes, fibroblast subsets, and glial cells, were associated with higher MCS scores, irrespective of the sampling time. Conversely, high expression of genes found primarily in epithelial cell populations, including goblet cells, BEST4+ enterocytes, enteroendocrine cells, and tuft cells, were associated with lower MCS scores (Figure 3A).

**Figure 3.**
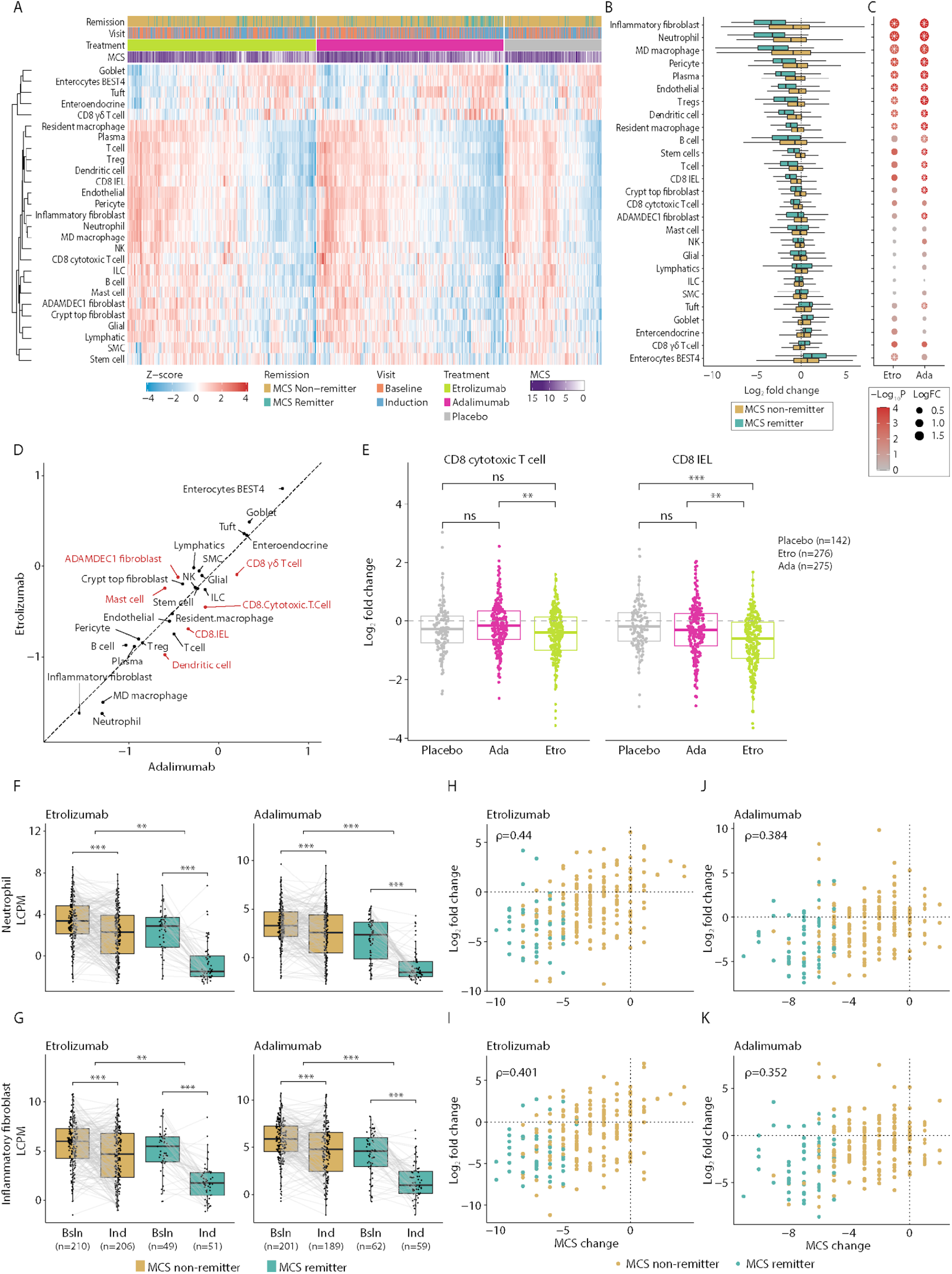
Neutrophils, monocyte-derived macrophages, and inflammatory fibroblasts are reduced by treatment, especially in remitters. **A.** Heatmap of signature scores for lineages defined in the single-cell atlas. Each row shows centered and scaled signature scores for a given population in individual samples from patients before or after treatment with etrolizumab, adalimumab, or placebo, as indicated. **B.** Effect of treatment on genes specific for individual cell populations in remitters or non-remitters after etrolizumab treatment (left panel). Boxes represent the log2 fold change between baseline and week 10 for either non-remitters or remitters. Cell populations are sorted by the magnitude of difference between longitudinal log2 fold change in remitters vs non-remitters. **C.** Significance of difference in log2 fold change between remitters and non-remitters after either etrolizumab or adalimumab treatment. Each point is colored by the p-value from comparing the effect in remitters and non-remitters, with the size showing the log2 difference between the fold changes. White stars indicate a >1.5 fold statistically significant difference in remitters vs non-remitter response at an FDR of 0.05. **D.** Effect of adalimumab (x-axis) or etrolizumab (y-axis) on individual cell type expression signatures. Each point represents an individual cell population, colored by those significantly different between etrolizumab and adalimumab (red) or not (black). **E.** Log2 fold changes of selected T cell populations after placebo, adalimumab, or etrolizumab treatment. Each point represents the difference between week 10 and baseline measurements of a cell type for either CD8+ IELs (left) or CD8+ cytotoxic T cells (right). Stars indicate nominal Wilcoxon signed-rank test p-values: ns: p > 0.01; *: p < 0.01; ** p < 0.001; *** p < 0.0001. **F.** Expression of neutrophil signature score before and after treatment with either etrolizumab or adalimumab. Each point represents a patient sample before (baseline) or after (induction, week 10) treatment. Samples collected from the same patient are linked by a line. **G.** Expression of inflammatory fibroblast signature score before and after treatment with either etrolizumab or adalimumab. Each point represents a patient sample before (baseline) or after (induction, week 10) treatment. Samples collected from the same patient are linked by a line. **H.** Comparison of log2 fold change in neutrophil gene expression signature (y-axis) with change in Mayo Clinic score (x-axis) after treatment with etrolizumab. Each point represents an individual patient, colored by remission status at week 10. **I.** Comparison of log2 fold change in inflammatory fibroblast gene expression signature (y-axis) with change in Mayo Clinic score (x-axis) after treatment with etrolizumab. Each point represents an individual patient, colored as in H. **J.** Comparison of log2 fold change in neutrophil gene expression signature (y-axis) with change in Mayo Clinic score (x-axis) after treatment with adalimumab. Each point represents an individual patient, colored as in H **K.** Comparison of log2 fold change in inflammatory fibroblast gene expression signature (y-axis) with change in Mayo Clinic score (x-axis) after treatment with adalimumab. Each point represents an individual patient, colored as in H. For panels F and G, stars indicate Benjamini-Hochberg corrected p-values: ns: p > 0.01; *: p < 0.01; ** p < 0.001; *** p < 0.0001.

Comparing changes in cell type-specific modules in remitters and non-remitters after ten weeks of induction treatment with etrolizumab demonstrated that the largest differences were found in neutrophils, MD macrophages, and inflammatory fibroblasts (Figure 3B). Consistent with these findings comparing inflamed and uninflamed samples in the single cell dataset, a significant increase in genes associated with BEST4+ enterocytes was observed after treatment, with more substantial increases found in remitters than non-remitters. The average change in cell-type-specific genes in patients treated with etrolizumab was inversely correlated with the differential abundance of these cell populations in inflamed vs uninflamed biopsies (Spearman correlation = −0.86, Figure S4B). Changes post-treatment appear largely treatment agnostic, with most populations showing similar behavior between etrolizumab and adalimumab treatment arms (Figure 3B, S4C, 3C). Notable exceptions include CD8+ T lymphocyte subsets, including IELs, CD8+ cytotoxic T cells and ɣδT cells, and dendritic cells, which were reduced to a greater extent following etrolizumab treatment, and ADAMDEC1+ fibroblasts and mast cells, which were reduced more following adalimumab treatment (Figure 3D-E). The cell specificity of the integrins was also observed to vary, with ITGAE, ITGB7, and ITGA4 showing higher specificity for T cell subsets, while ITGB1 was expressed on many stromal cells. ITGB7 and ITGA4 were additionally expressed on dendritic cells, resident macrophages, and plasma and B cell subsets (Figure S4D). Comparing relative changes across neutrophil and inflammatory fibroblast-associated genes, greater reductions were observed in patients who achieved MCS remission (Figure 3F-G), while non-remitters showed less reduction. Furthermore, reductions in neutrophil and inflammatory fibroblast-associated genes were correlated with overall changes in MCS irrespective of remission status or biologic treatment (Figure 3H-K, Figure S4E-H, Spearman rho = 0.44 for neutrophils, rho = 0.40 for inflammatory fibroblasts). Supporting the role of neutrophil subsets and inflammatory fibroblast as crucial cell types for MCS remission, previous anti-TNF resistance modules were expressed at the highest levels by both cell types (12) (Figure S5A).

### Heterogeneity of neutrophils in colonic tissue of UC patients

Given the notable increase in neutrophils within inflamed tissue relative to uninflamed tissue and their association with persistent disease despite etrolizumab and adalimumab treatment, it was of interest to further characterize these cells to understand the heterogeneity of these populations in inflammatory tissue such as UC. Four major subsets of neutrophils were identified within inflamed and uninflamed biopsies, along with one subset of neutrophils that could not be further classified due to low sequencing depth (Figure 4A, Figure S6A). All subsets were confirmed to express neutrophil lineage markers, including FCGR3B, VNN2, CXCR2, and PROK2 while lacking expression of marker genes indicative of closely related myeloid cells, such as VCAN and CD300E expressed in MD macrophages, HLA-DR in macrophages and dendritic cells, or CLC in eosinophils (Figure 4B). Specific markers such as CXCR4, OSM, MX1, and PADI4 were used to classify these populations for each neutrophil subset. MX1hi neutrophils co-expressed several markers consistent with interferon signaling, including MX1 itself, IFI6, IFIT2, and IFIT3 (Figure 4C). OSMhi neutrophils expressed high levels of inflammatory cytokines and chemokines, including OSM, IL1B, and CXCL1 (Figure 4C, Table S5). PADI4hi neutrophils expressed high levels of several membrane-bound proteases, including MMP9, MMP25, and MME (Figure 4C, Table S5). Lastly, CXCR4hi neutrophils produced a unique set of cytokines: CCL3, CCL4, VEGFA, and CSF1 (Figure 4C). All four subsets of neutrophils had significantly higher abundance in inflamed colonic tissue in comparison to uninflamed tissue (Figure 4D, Figure S6B). A smaller dataset of neutrophils from colonic biopsies has been reported with the identification of 3 neutrophil subsets, N1, N2, and N3 (18), which exhibited similar expression patterns to PADI4/OSM, CXCR4, and MX1 neutrophils, respectively (Figure S6C).

**Figure 4.**
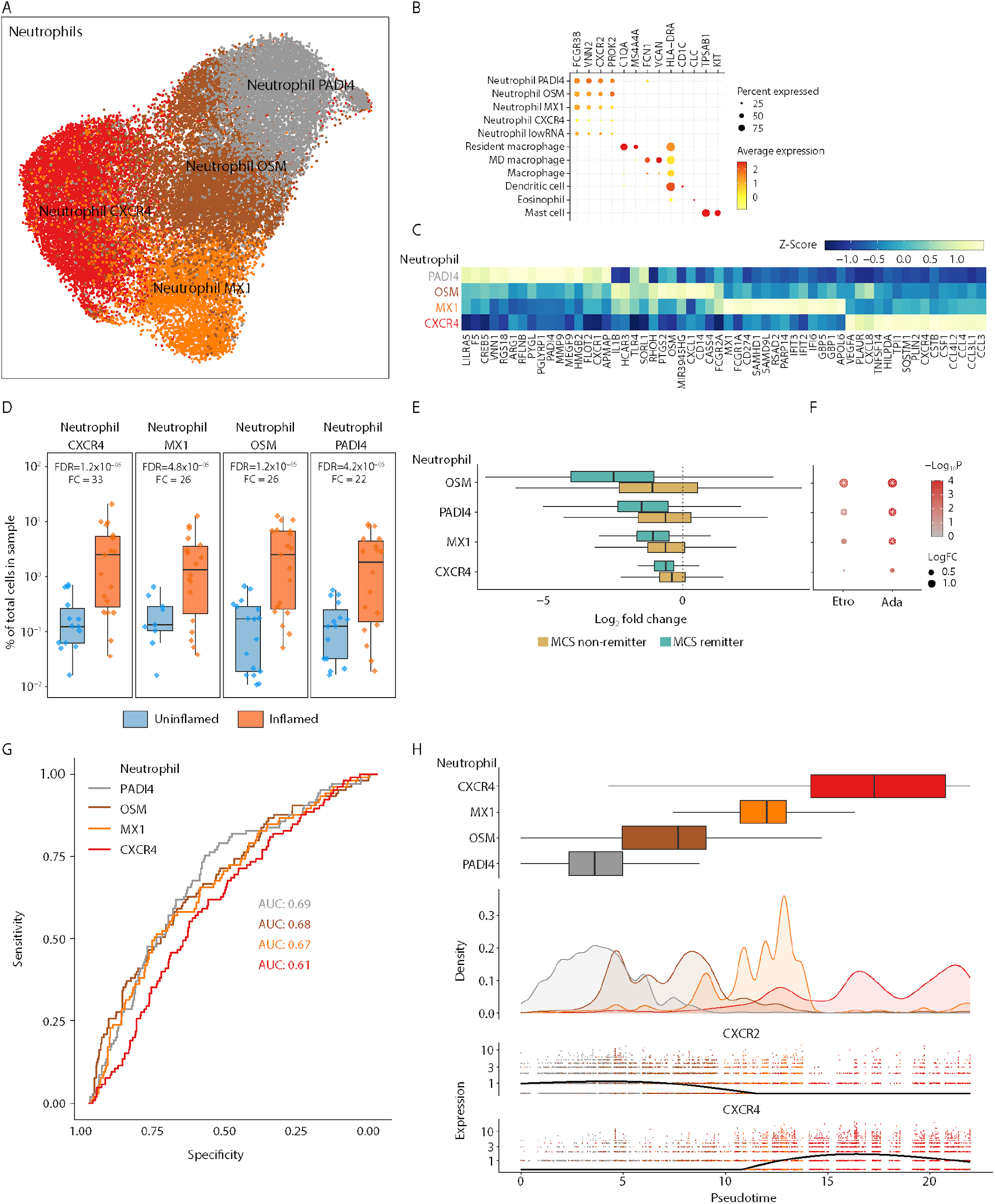
Heterogeneity of neutrophils in UC tissue biopsies. **A.** UMAP projection of neutrophil subsets. Each point represents a cell, colored by the respective subset it was classified as. **B.** Dotplot of marker genes for myeloid subsets. Each point represents the expression of a gene in a cell population. The size of the point indicates the proportion of cells expressing the gene, and the color indicates the centered and scaled expression level. **C.** Heat map of marker genes for each neutrophil subset. Colors show centered and scaled gene expression values across four subsets of neutrophils. **D.** Differential abundance of neutrophils across inflamed and uninflamed tissue. Each point represents a sample. The y-axis shows the fraction of each neutrophil subset relative to the total number of cells in that sample. Fold changes and FDR values were determined as described in the methods. Boxes represent the lower and upper quartiles, with the center line representing the median. Whiskers extend to the most extreme point no more than 1.5x the IQR from the box boundaries. **E.** Log2 fold changes of gene expression signatures from each neutrophil subset after etrolizumab treatment, split by remission status at week 10. Cell populations are sorted by the magnitude of difference between longitudinal log2 fold change in remitters vs non-remitters. **F.** Significance of difference in log2 fold change between remitters and non-remitters after either etrolizumab or adalimumab treatment. Each point is colored by the p-value from comparing the effect in remitters and non-remitters, with the size showing the log2 difference between the fold changes. White stars indicate a >1.5 fold statistically significant difference in remitters vs non-remitter response at an FDR of 0.05. **G.** ROC curves for the association of longitudinal changes in neutrophil subsets with remission **H.** Pseudotime inference for neutrophil subsets. The x-axis represents the ordering of cells through pseudotime. Boxplots show the relative placement of neutrophil subsets across pseudotime, with density plots showing the total distribution of cells. Normalized expression values for CXCR2 or CXCR4 are shown across pseudotime, with the y-axis representing log2 normalized counts, each point representing a cell positioned in pseudotime and colored by the subset it belongs to.

Comparing these gene sets representative of these four neutrophil subsets in pooled HIBISCUS datasets, the most extensive changes post-treatment were observed in genes expressed by OSMhi and PADI4hi neutrophil populations, which also showed significant differences in treatment effect between remitters and non-remitters (Figure 4E). These gene sets were specific to each neutrophil subset, and this effect was treatment agnostic, as the same pattern was observed after etrolizumab and adalimumab treatment (Figure S6D, Table S6, Figure 4F). In determining if these neutrophil signatures (Table S6) could serve as predictors of remission, a similar pattern of marked reduction of PADI4hi and OSMhi-specific genes was more closely associated with remission, suggesting that these neutrophil subsets were related to residual disease (Figure 4G). Pre-treatment levels of these neutrophil cell subsets only showed modest performance as prognostic biomarkers (Figure S6E).

Previous reports have indicated that neutrophils regulate the expression of CXCR2 and CXCR4 throughout their lifecycle (22). Consistent with this finding, a gradient of CXCR2 and CXCR4 expression across the four subsets of neutrophils was observed, suggesting that these subsets might represent points along a cell differentiation trajectory (Figure S6F). Pseudotime inference was used to create a trajectory through our neutrophil single-cell data, and consistent with this model, a gradient of decreasing CXCR2 and increasing CXCR4 expression was observed through pseudotime (Figure 4H). The four subsets we identified fell into broad regions of the pseudotime axis, with the trajectory running from PADI4hi, then OSMhi, through MX1hi, and ending with the CXCR4hi state (Figure 4H). Consistent with this proposed pseudotime model, a subpopulation of MMP9 (gelatinase B) and bactericidal permeability-inducing protein (BPI) expressing cells within the PADI4hi subset was identified, suggesting the presence of immature neutrophils (Figure S6G) (23, 24). Cytokines/chemokines such as CCL3, CCL4, and VEGFA were expressed by more mature neutrophils in the inferred pseudotime trajectory (Table S7, Figure S7A).

### Rewiring of cell-cell communication in neutrophil subsets in the inflamed tissue

Given the increased abundance of neutrophil subsets in inflamed tissue and previously described association with inflammatory fibroblasts (12), elevations in inflamed tissue, and modulation by etrolizumab and adalimumab, we analyzed cell-cell communication networks to characterize predicted interactions between neutrophils states and stromal cells. The most substantial predicted ligand-receptor interactions were between all neutrophil subsets with multiple stromal populations, especially in inflamed tissues compared to uninflamed tissue, with both PADI4hi and OSMhi neutrophil subsets showing the most significant difference in interaction potential (Figure 5A, S8A). Meaningful interactions within signaling pathways from inflammatory fibroblasts to all neutrophil subsets were also identified (Figure 5B, S8B). The most chemokine interactions were from inflammatory fibroblasts to PADI4hi, OSMhi, and MX1hi neutrophil subsets via CXCL1/2/3/5/6/8 expressed by fibroblasts and CXCR1/2 expressed on neutrophils (Figure 5C). In addition, interactions were predicted between crypt-bottom fibroblasts, CD36 endothelial cells, and CXCL12+ endothelial cells with CXCR4hi and MX1hi neutrophils via CXCL12 expressed by stromal cells and CXCR4 expressed by neutrophils (Figure 5C). Among all ligand-receptor interactions mentioned, only CXCR4:CXCL12 interactions were present between these stromal cells and CXCR4hi neutrophils in uninflamed tissue, as were CXCL1:CXCR2 interactions between inflammatory fibroblasts and PADI4hi, OSMhi and MX1hi neutrophils, albeit in both cases the signaling strength was reduced in uninflamed tissue (Figure S8B, S8C).

**Figure 5.**
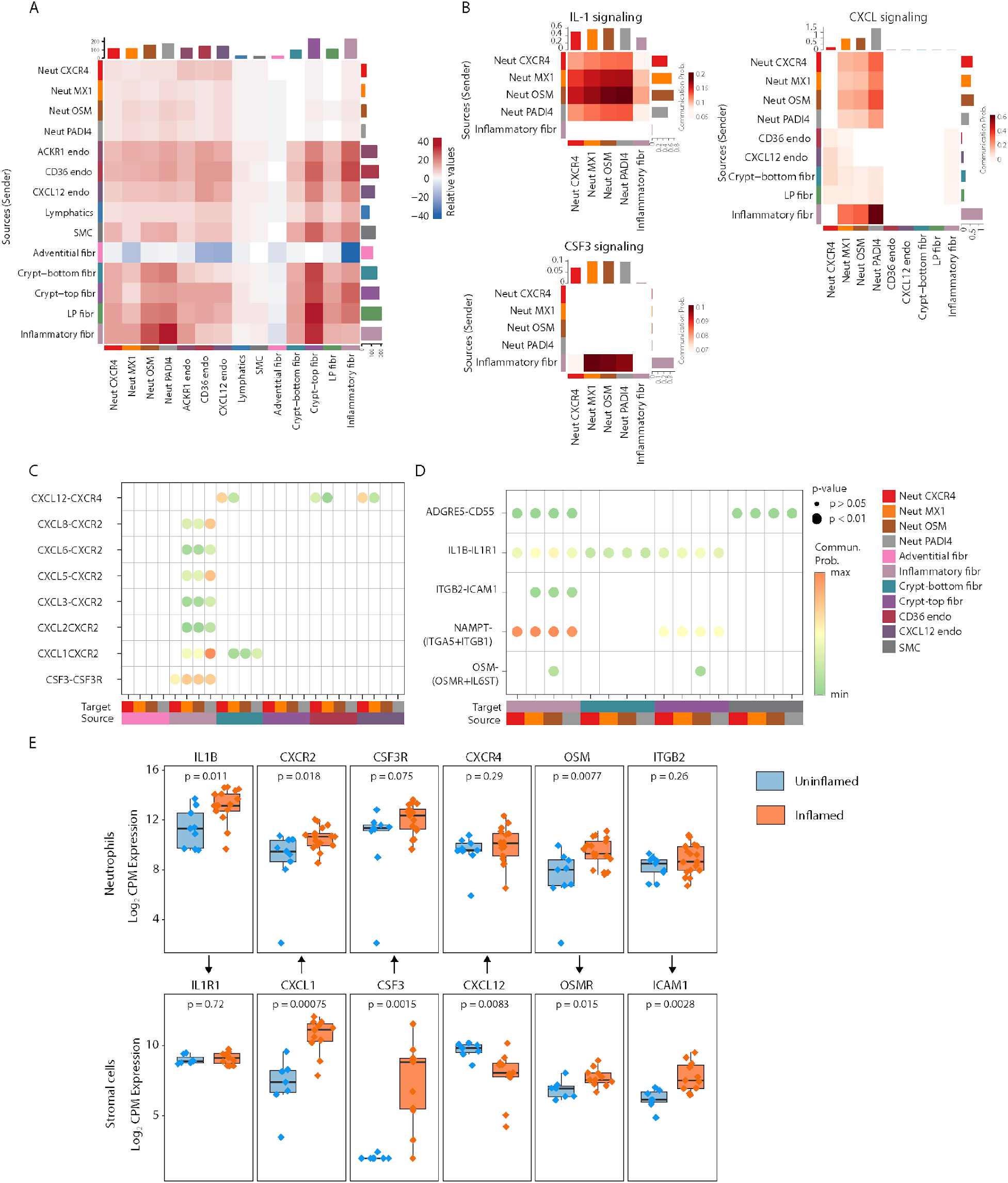
Neutrophil interaction rewiring in UC inflamed tissue. **A.** Heatmap of the differential number of inferred interactions between inflamed and uninflamed tissue among neutrophil and stromal subpopulations. **B.** Heatmap of signaling pathways CXCL, CSF3, IL1 showing significant communications between neutrophil subpopulations and inflammatory fibroblasts in inflamed tissue. The color in the heatmap is proportional to the overall communication probability between each cell type. **C.** Selected significant ligand-receptors as incoming signaling interactions to neutrophil subpopulations from fibroblasts and endothelial subsets in the inflamed tissue. **D.** Significant ligand-receptors were selected as outgoing signaling interactions between neutrophil subpopulations, fibroblasts, and endothelial subsets in the inflamed tissue. **E.** Pseudo-bulk expression levels of ligands and receptors in the neutrophil population. Expression values are shown on the y-axis as log2 normalized counts per million (CPM). **F.** Pseudo-bulk expression levels of ligands and receptors in the stromal population. Expression values are in log2 normalized counts per million (CPM). Based on the Wilcoxon test, P-values are reported for significant change in expression between the inflamed (orange) and uninflamed (blue) tissue.

Furthermore, significant IL-1 signaling was evident from all neutrophil populations to inflammatory fibroblasts, with more robust signaling from PADI4hi and OSMhi neutrophils (Figure 1E, 5B, 5D). This interaction was also found within uninflamed colonic tissues (Figure S8D, S8B). Conversely, we predict CSF3 signaling from inflammatory fibroblasts with all neutrophil populations through CSF3:CSF3R (Figure 5B, 5C). OSM: OSMR signaling was also identified as a strong interaction between neutrophils, specific to OSMhi neutrophils, and inflammatory and crypt-top fibroblasts (Figure 5D). Beyond cytokines and chemokines, other notable interactions were observed acting on receptors on inflammatory fibroblasts, including NAMPT: ITGA5/ITGB1, ITGB2:ICAM, ADGRE5:CD55. The increased expression of receptors and ligands specific to the neutrophils and fibroblast populations between inflamed and uninflamed tissue supports the predicted increased crosstalk between these populations (Figure 5E-F).

## Discussion

Ulcerative colitis is an inflammatory immune condition resulting from genetic predisposition, environmental triggers leading to alterations in the gut microbiome, and via dysregulation of the innate and adaptive immunity resulting in damage to the intestinal epithelial barrier (1–3). Immune infiltrates contributing to disease are characterized by adaptive and innate immune cell populations, including T and B lymphocytes, plasmablasts, and diverse myeloid populations (11, 13, 14). For example, IL17-expressing T cell subsets are reported to expand in the inflamed gut of UC patients, contributing to the TH17-driven pathobiology and tissue damage (11). The GIMATS module reported by Martin et al., comprising activated DCs, inflammatory macrophages, activated T cells, IgG plasma cells, activated fibroblasts, and endothelial cells, as well as myeloid infiltration into the colonic epithelium reported by Jha et al. were shown to be associated with non-response to anti-TNFs and more severe disease (13, 14). These various cellular compartments work in concert to execute immune-mediated processes that include but are not limited to direct cytotoxicity against epithelial cells, aberrant B cell-mediated antibody production with likely resultant macrophage activation, resistance to regulatory T cells, and, importantly, ulceration caused by infiltrating neutrophils (13, 25–31). In the non-immune compartment in affected UC tissue, it has been reported that there is a greater abundance of inflammatory fibroblasts that, while not necessarily leading to strictures described in Crohn’s disease, may sustain continued recruitment of immune cell populations such as myeloid cells via cytokine/chemokine signaling (12, 14, 15). Breaking this cycle of inflammation has been successful in some patient populations with anti-TNFs, anti-integrins, JAK inhibitors, and, more recently, IL-23 and TL1A inhibitors. However, patterns of non-response across these agents appear consistent in that the presence of inflammatory cells and inflammatory pathways are not fully normalized relative to patients in remission (10–17, 32, 33).

Etrolizumab Phase 3 studies did not meet the desired endpoint in all studies (9, 34–36); however, they provided sufficient numbers of well-characterized patients with differential clinical outcomes and biomarker assessments, allowing for a rigorous evaluation of molecular changes in colonic tissues assessed before and after induction treatment. This report shares results from pooled HIBISCUS I & II studies. These were identical placebo-controlled studies that compared the clinical efficacy of etrolizumab and adalimumab, conducted in biologic-naive patients to minimize heterogeneity introduced by prior treatment. Biomarker data from both trials was aggregated to allow a more rigorous assessment of patients treated with etrolizumab, adalimumab, or placebo by treatment outcome. While not shown, molecular changes described in the colonic tissue in HIBISCUS were representative of those observed in other Phase 3 UC studies of etrolizumab, including LAUREL, GARDENIA, and HICKORY.

Overall, we observed significant reductions in genes expressed by memory T and B cell subsets, including genes that encode alpha 4, alpha E, and beta 7 integrins that form heterodimeric complexes and allow for mucosal tissue homing. Despite the significant reductions in ITGA4 and ITGB7, there was minimal to no change in ITGB1, potentially due to expression in other cellular populations (Figure S4D). Similar observations were made in post-treatment biopsies from a Phase 3 etrolizumab study in Crohn’s disease (7). Although these findings align with the proposed dual action of etrolizumab, similar results were seen with adalimumab treatment, suggesting the changes are due to reduced inflammatory infiltrate following a decrease in disease severity. A unique pharmacodynamic-specific effect observed was reduced expression of ITGAE and IEL-specific genes only under etrolizumab treatment, implying reduced retention or increased egress of aEb7+ expressing lymphocytes. This insight is backed by findings from the EUCALYPTUS and mechanistic in vivo studies (6, 7). Analyzing colonic samples earlier in the treatment cycle could give more precise insights into direct mechanistic and pharmacodynamic effects.

Both etrolizumab and adalimumab treatments caused substantial changes in the transcriptional profiles of resident colonic cells, decreasing inflammation-associated genes and slightly increasing epithelial cell-related genes. The degree of these alterations correlated with changes in MCS or remission status. Interestingly, alterations in myeloid lineage and inflammatory fibroblast gene expression were closely associated with MCS changes. Previous studies indicated that non-response to TNF blocking agents is most strongly linked to myeloid-expressed genes, including OSM and IL1B, among others (12, 37–39). Our data confirmed this finding (Figure S5A) and extended this conclusion to other MOAs, such as anti-integrins, suggesting that the association observed indicates a measure of disease state or disease resolution rather than reflecting specific mechanistic changes. Based on these results, the concept of disease resolution in ulcerative colitis might evolve beyond endoscopic and histological healing to include restoration of specific molecular pathways altered in disease pathogenesis (40, 41).

Despite the relevance of myeloid cells, single-cell data on specific subsets, particularly neutrophils, have been limited due to technical challenges associated with cell viability and low transcript levels. Herein, we described distinct myeloid populations, identified via the generation of a single-cell RNA-sequencing dataset which included over 400K cells, including >30K neutrophils, allowing for detailed characterization of myeloid populations, including monocyte, macrophage, dendritic cell, and neutrophil sub-populations, to a greater extent than was previously possible. As expected, we observed an increased abundance of neutrophils, MD macrophages, and inflammatory fibroblasts in inflamed tissue. We saw a significant reduction in BEST4+ enterocytes. The bulk transcriptomic data from the HIBISCUS trials also reflected this inverse relationship between immune and epithelial cells. Referencing our single-cell dataset to identify cell lineage and cell subset markers, when applied to the HIBISCUS dataset, revealed significant reductions in neutrophils, MD macrophages, and inflammatory fibroblasts with both etrolizumab and adalimumab induction treatment, supporting the role of these cells in disease pathobiology. These cells have also been described to be perturbed even within the epithelial layer as being disrupted and evidence of resident ɣδT cells, TH17 cells, and inflammatory myeloid cells (13).

We identified four sub-populations of neutrophils that exhibited distinctive gene expression profiles and could be discriminated using CXCR4, OSM, MX1, and PADI4 expression. The diversity of differentially expressed genes across these subsets of cells highlights the variety of processes that neutrophils mediate. One additional subset of neutrophils with low RNA abundance was observed; however, it was unclear if this was a technical artifact or if this population has some biological relevance, and excluded it for further analyses. In our analyses, we found all four neutrophil subsets present in both inflamed and uninflamed biopsies, with no difference in relative abundance across the subsets. In HIBISCUS, not all neutrophil subsets contributed equally to the likelihood of remission; we observed that reductions in OSMhi and PADI4hi subsets appeared to have the closest association with clinical remission with no association between decrease in CXCR4hi neutrophils and remission status.

We hypothesized that the individual states observed in our data represent points along the differentiation trajectory for neutrophils as they enter the tissue. In line with this hypothesis, pseudotime inference indicated a potential cell state trajectory from the least mature to the most mature neutrophil population, aligning with the CXCR2:CXCR4 axis (22, 42, 43). Accordingly, gene expression across colonic UC neutrophil subsets overlaps with tissue and blood neutrophil transcriptional signatures from previous studies that demonstrated differentiating cell state trajectories in rodents (42). Consistent with this model, we found the highest expression of immature neutrophil markers, including MMP9 (gelatinase B) and BPI (23, 24) in PADI4hi cells, with reduced expression along our pseudotime trajectory.

With differences in cell state, we predicted some functional differences that could be inferred via studies of potential ligand-receptor interactions between neutrophils and other resident cells, such as stromal cells. Several signaling pathways were enriched at the interface between OSMhi and PADI4hi neutrophils with inflammatory fibroblasts, including OSM: OSMR and IL1B: IL1R. Friedrich et al. also identified IL-1b-driven interactions between neutrophils and fibroblast subsets, independent of TNF-a, suggesting a key mechanism for resistance to anti-TNF therapies (12). In addition, we found an interaction between crypt-bottom fibroblasts and more mature neutrophils via CXCL12:CXCR4 signaling that persisted in both inflamed and uninflamed conditions. Within the stromal compartment, inflammatory fibroblasts produce CXCL1/2/3/8, key ligands associated with neutrophils homing, through their interaction with CXCR1/2. Together with the expression of CSF3 as a crucial regulator of neutrophil production and increased lifespan, the data highlights a potential role for neutrophils’ immune response organization via cytokine signaling in the stromal compartment. Previous reports highlighted the role of CXCR4:CXCL12 signaling in IBD through homing of peripheral blood and lamina propria T cells, and migration of B cells from Peyer’s patches (44–46) and a role for CXCL12 in both the inflamed state and homeostasis (44). How each of these immune cell compartments responds to CXCL12 signaling and under what conditions bears further study. The relevance of CXCR4hi neutrophils to IBD pathobiology is unclear from our results since, despite making up a significant fraction of total neutrophils, they are less associated with disease state than less mature populations. CXCR4 has been described as a mechanism that facilitates the return of aging neutrophils to the bone marrow and retention within inflamed tissues, which may be reflected in this dataset and others. These findings highlight a potential role for neutrophils in organizing the acute immune response via cytokine signaling in the stromal compartment.

Neutrophils have been appreciated as a locus for therapeutic intervention in IBD and other inflammatory diseases and are linked to therapeutic non-response (17, 47). In mouse colitis models, CXCR2 blockade reduced neutrophil function and inflammatory cytokine production and reduced pathology (48–50). Several molecules that block CXCR2 signaling have been tested as therapeutic interventions across many diseases, but little information has been reported in inflammatory bowel disease (51). Likewise, little information exists about targeting IL-1b signaling in IBD outside of very early onset IBD (52, 53). While neutrophils are likely drivers of the chronic pathology of ulcerative colitis, neutrophils are necessary to maintain host integrity at sites of infection and minimize signs of inflammation. When neutrophils are compromised, there is a risk of a lack of mucosal defense and potential for serious clinical manifestations; therefore, direct targeting of neutrophils is undesirable, especially in chronic diseases.

These results characterizing the phenotype and potential interactions between neutrophils and other resident cells may lead to alternative approaches for therapeutic intervention. Given the consistent finding of myeloid cells in non-response with several agents, we hypothesize that interference of cell-cell interactions or response to local products necessary for neutrophil infiltration, retention, or activation may be necessary to achieve greater rates of clinical remission in UC patients with moderate to severe disease.

## Materials and Methods

### Clinical trial bulk RNA-sequencing

#### Study design and patients

Available colonic biopsies were assessed from TNF inhibitor-naive adult patients with moderate-to-severe UC enrolled in two identically designed, randomized, double-blind, placebo-controlled Phase 3 studies, Hibiscus I and Hibiscus II (9). As previously reported, randomized patients received subcutaneous doses of either placebo (n=137), etrolizumab (n=259), or adalimumab (n=263). We compared samples at baseline (pre-dose) and at the end of induction at week 10. The primary endpoint was the induction of remission at week 10, defined as a Mayo Clinic Score (MCS) of 2 or lower, with individual subscores of 1 or lower and rectal bleeding subscore of 0. Additional study and patient details can be found in the published clinical report (9).

#### Biopsy collection

Colonic biopsy pairs were collected from the most inflamed region (20–40 cm from the anal verge) of patients undergoing a colonoscopy procedure, and biopsies were immediately stored in RNA*later* solution (Cat.#AM7022, Invitrogen).

#### Biopsy processing

Frozen colonic tissue pinch biopsies from trial subjects, stored in RNAlater (Qiagen) tubes, were thawed and placed into individual wells of a PowerBead Block (Qiagen) containing 0.1 mm glass beads and 450 µl RLT buffer (Qiagen) plus 2-beta mercaptoethanol (Sigma). Blocks were then sealed and homogenized using TissueLyser (Qiagen) at 25 hertz for 2 minutes. Blocks were then rotated, and an additional homogenization step was performed. The resulting lysate was removed from the block after centrifugation, and RNA/DNA was simultaneously isolated with an All-Prep 96 HT kit (Qiagen) according to the manufacturer’s instructions.

#### Bulk RNA sequencing

As detailed above, purified RNA isolated from colonic pinch biopsies were assessed for quantity and purity using Qubit and integrity with Agilent Tapestation. RNA of high quality then proceeded to library preparation with the Illumina TruSeq mRNA stranded kit. The resulting libraries were then evaluated for quantity (Quant-It) and fragment size determination (Agilent Tapestation). Successfully generated libraries were loaded onto a NovaSeq 6000 instrument and sequenced with a total read length of 1x50bp, each sample reaching at least 27 million single-end reads. Once sequencing was complete, the quality of the reads was assessed for clusters passing the filter, Q30 scores, error rate, cluster density, and read distribution before proceeding further for downstream workflow analysis.

#### RNA-seq alignment and differential expression analysis

RNA-seq data was processed using HTSeqGenie (54). Reads were filtered and aligned to Genome Reference Consortium Human Build 38 (GRCh38) using GSNAP (55), and uniquely mapping reads to gene models present in the GENCODE basic annotation set (v. 27) were used for generating the gene count matrix and downstream analyses. Additional analysis and visualizations were performed in R version 4.2.0 (https://www.r-project.org/). The EdgR package was used for preprocessing, including filtering of low-count genes, trimmed mean of m-values (TMM) normalization, and log counts per million (lcpm) transformation of counts. Differential gene expression analysis was performed using the voom-limma framework with the subject introduced as a blocking factor.

### Single-cell RNA-seq atlas

#### Patient and tissue sample collection

Subjects were enrolled in the Stanford Inflammatory Bowel Disease (IBD) Registry used for all ulcerative colitis (UC) patients and healthy controls under Institutional Review Board (IRB) Protocol 52317 at Stanford University School of Medicine. Informed consent was obtained from all patients in accordance with the respective protocol, and sequencing data storage and publication plans were approved by the Stanford IRB. UC patients were included based on having a clinical diagnosis of ulcerative colitis and observed to have active disease via macroscopic assessment from a physician during an endoscopy as part of routine clinical care. Biopsies were obtained during endoscopy, using biopsy cold forceps used in standard care. The endoscopist visually evaluated the presence or absence of inflammation at the time of collection. We used the Mayo Endoscopic Score (MES) to ensure this evaluation was consistent across endoscopists. UC patients without endoscopic inflammation were denoted with a MES of 0, whereas those with active inflammation were denoted as MES 1 (mild inflammation), MES 2 (moderate inflammation), and MES 3 (severe inflammation). Four to six biopsies were obtained from inflamed and uninflamed colonic segments from 20 UC patients. Biopsy bites were immediately placed into Advanced DMEM F-12 media (Gibco, catalog number 11320033), placed on wet ice for transport, and immediately processed fresh for single-cell experiments. Only fresh biopsies were included in this study.

#### Epithelial layer dissociation

On arrival, biopsy bites were washed twice in cold PBS. Tissue was then added to 5 mL of an EDTA-enriched epithelial dissociation medium [HBSS without Ca^2+^Mg^2+^ (Life Technologies #88284), HEPES 10 mM (Life Technologies #15630106), EDTA 10 mM (Life Technologies #AM9260G], 100 U/ml penicillin (ThermoFisher #15140-122), 100 mg/mL streptomycin (ThermoFisher, #15140-122), FBS 2% + freshly supplemented with 100 µL of 0.5M EDTA] and placed in a rotisserie incubator at 37°C for 15 minutes. Following incubation, the tissue rested on ice for 10 minutes and was shaken vigorously for 15 seconds. Remnant tissue was removed and placed on ice-cold PBS for downstream dissociation of the lamina propria layer. The remaining supernatant containing the epithelial layer was spun at 800 g for 5 minutes, resuspended in 1 mL of epithelial dissociation medium, and transferred to a 1.5 mL Eppendorf tube. The epithelial solution was spun down at 1000 g for 2 minutes and then resuspended in 1 mL of TrypLE express enzyme (ThermoFisher #12604013) for 5 minutes at 37°C, followed by gentle trituration with a P1000 pipette. Epithelial cells were spun down at 1000 g for 2 minutes, resuspended in 200 µL of epithelial cell solution, and placed on ice for 3 minutes before triturating with a P1000 pipette. The solution was filtered into a new Eppendorf tube through a 100 µM cell strainer (Falcon/VWR #21008-949). The epithelial cells were then spun down at 1000 g for 5 minutes, resuspended in 1 mL of PBS with 0.04% BSA, and kept on ice until single-cell experiments.

#### Lamina propria layer dissociation

Tissue saved on ice from epithelial layer digestion was transferred into 5 mL of digestion medium [RPMI 1640, 100 U/ml penicillin (ThermoFisher #15140-122), 100 mg/mL streptomycin (ThermoFisher #15140-122), 50 mg/mL gentamicin (ThermoFisher #15750-060)], FBS 2%, HEPES 10 mM (Life Technologies #15630106), and freshly supplemented with 100 µg/mL of Liberase TM (Roche #5401127001) and DNase I 100 µg/mL (Sigma-Aldrich # LS002060) and placed in a rotisserie incubator at 37°C for 30 minutes. After incubation, the enzymatic dissociation was quenched by adding 1 mL of 100% FBS + and 80 µL of 0.5M EDTA and placed on ice for five minutes. The solution was filtered through a 100 µM filter into a new 50 mL conical tube and then transferred to a new 15 mL Falcon tube. The solution was spun down at 1000 g for 5 minutes, resuspended in 1 mL of PBS with 0.04% BSA, and then transferred to a new 1.5 mL Eppendorf tube. The cells were then spun down at 1000 g for 5 min and resuspended in 1 mL of 1X PBS with 0.04% BSA (Invitrogen #AM2616).

#### Single-cell RNA-sequencing

Epithelial and lamina propria single-cell suspensions were counted and, if necessary, diluted to a concentration of 200–500 cells per µL. About 40,000–60,000 cells (1:1 ratio of epithelial to lamina propria cells) from each sample were loaded into the BD Rhapsody™ Cartridge (BD Biosciences; Cat. No. 633733). According to the manufacturer’s instructions, single-cell capture, barcoding, lysis, and cDNA synthesis were performed with the BD Rhapsody™ Express Single-Cell Analysis System.

According to the manufacturer’s instructions, whole transcriptome analysis (WTA) libraries were indexed and prepared using the BD Rhapsody™ WTA Amplification Kit (BD Biosciences, Cat. No. 633801).

Libraries were sequenced on Illumina MiSeq, HiSeq 2500, NextSeq 2000, and NovaSeq 6000 (Illumina) with the following configuration: Read 1, 75bp, i7 index: 8bp, i5 index: none, Read 2, 75bp.

#### Individual library preprocessing

To process samples aligned to a reference genome, we obtained raw gene cell count matrices for each sequenced sample from the Seven Bridges pipeline and processed them using the Seurat package V4 (56). We removed low-quality cells from our analysis by excluding cells if we detected less than one hundred genes or more than twenty-thousand RNA counts per cell. We followed the standard Seurat pipeline by performing log normalization of our counts and scaling and performing linear dimension reduction analysis using the top two thousand variable features for Principal Component Analysis (PCA). We then found K nearest neighbors and detected communities using a Louvain Jaccard clustering algorithm that used euclidean distance to identify communities using our PCAs to create a Uniform Manifold Approximation and Projection (UMAP) reduction. As part of the inclusion criteria for each sample, we confirmed that each library had an appropriate representation of expected cell types in the gut. For example, we confirmed that we had a representation of immune and epithelial cells before including them in downstream analysis. One inflamed and one uninflamed sample did not pass quality control and was omitted, resulting in 38 overall samples, with 18 pairs of inflamed and uninflamed samples across 20 patients.

#### Merging and integration

All sequenced libraries for paired inflamed and uninflamed biopsies were individually processed and sequenced, which can introduce batch-specific effects across different samples. To reduce technical and biological batch variability, we first merged all of our samples and then integrated them using the Harmony package (57). Harmony provides a robust approach that projects cells into a shared embedding space based on cell type. Specifically, we used the top 50 dimensions from our PCA reduction analysis to integrate our merged biological samples.

#### Annotation at the single-cell level using SCimilarity

To annotate our cells, we first ran our integrated dataset through SCimilarity (58). SCimilarity leverages existing publicly available scRNA-seq atlases to train a deep-learning model to identify single cells. This model can then be applied to novel instances of single cell data where it calculates the distance between each individual cell and cells within the model. Specifically, we used the “ca.annotate_dataset()” function to annotate cells at the individual level with the “2023_01_rep0” model. After annotating cells at the individual level, we grouped each assigned cell annotation to one of the following lineages: T cells, B cells, epithelial cells, stromal cells, and myeloid cells. Each lineage was separated into an individual dataset for downstream analysis and manual annotation.

#### Doublet and low-quality cell removal

We iteratively removed doublets to improve community detection as we annotated each lineage. We created separate objects based on their cell lineage to remove doublets, reharmonized, and reclustered. Notably, it was essential to reharmonize each time as it improved gene detection within each cluster. We classified doublets as cells that coexpressed canonical cell type markers across different lineages. For example, we removed clusters that coexpressed EPCAM, a canonical epithelial cell marker, and CD3E, a canonical T cell marker, from our T cell or epithelial cell object. We repeated the above process until we could not detect doublet clusters within each cell lineage.

Since epithelial cells often undergo apoptosis during library preparation, this lineage tended to have much higher levels of mitochondrial reads than other lineages. Therefore, to remove low-quality/apoptotic cells from the epithelial lineage, we excluded cells that contained more than seventy percent of their reads from mitochondria. This drastically reduced the number of doublets and low-quality cells present within the lineage and allowed for more accurate community detection.

#### Annotation of cell lineages

After removing doublets and low-quality cells from each lineage, we harmonized and clustered each lineage at varying degrees of granularity. We annotated each lineage at three different levels of granularity, with level 1 being the least granular and level 3 being the most granular. Cells were manually annotated using existing IBD atlases as references (11, 14, 59). We used marker genes’ expression in each cluster listed in our cell type marker table (Table S8) to assign identities.

Stromal cells were annotated as endothelial cells (PECAM1), fibroblasts (ADAMDEC1, ABCA8, LUM, DCN, POSTN), smooth muscle cells (MYH11, NPNT, HHIP), glial cells (ALDH1A1, S100B) and pericytes (RGS5, NDUFA4L2). Endothelial cells were further clustered and subdivided into ACKR1 endothelial cells (ACKR1), CD36 endothelial cells (CD36), CXCL12 endothelial cells (CXCL12), and lymphatics (LYVE1, CCL21). Fibroblasts were also clustered again and subdivided to adventitial fibroblasts (CD34, PI16, SFRP2), crypt-bottom fibroblasts (WNT2B, RSPO3), lamina propria (LP) fibroblasts (ADAMDEC1, ABCA8, CCL2), crypt-top fibroblasts (PDGFRA, F3, WNT5A, WNT5B) and inflammatory fibroblasts (IL11, IL13RA2, CHI3L1). Additionally, we observed a subset of crypt-top fibroblasts in activated states (PDPN).

Epithelial cells (EPCAM) were annotated as Enterocytes (CA1, CA2, SLC26A2/3, AQP8), Stem cycling (PCNA, LGR5, SMOC2, OLFM4), Goblets (MUC2, TFF1), Enteroendocrine (CHGA), and Tuft cells (SH2D6). BEST4+ Enterocytes (BEST4) were then annotated as a subset of enterocytes. Stem cycling cells were clustered again and divided into Stem cells (PCNA, LGR5) and Transit amplifying (TOP2A).

Myeloid cells were annotated as granulocytes (FCGR3B), mast cells (TPSAB1), red blood cells (HBB), or macrophages/DCs (HLA-DRA). Granulocytes were divided into neutrophils (FCGR3B) and eosinophils (CLC, HLA-DRA). Macrophages/DCs were further divided into resident macrophages (C1QA, C1QB), MD macrophages (VCAN, FCN1), or dendritic cells (CD1C). Dendritic cells were further classified as plasmacytoid DCs (IRF7, TNFRSF21), tertiary lymphoid structure-associated DCs (CCR7, CCL19), conventional DC1 (SLAMF8, CLEC9A), conventional DC2 (CD14, CLEC10A), or myeloid DCs (CD1A, ITGAE). While performing annotation, if a cluster of cells expressed multiple lineage-specific markers at high levels, those cells were marked as doublets and excluded from further analysis.

Assigned cell identities were then independently validated by at least one additional researcher to ensure proper annotation. After annotating cell lineage objects, all cell lineage objects were merged together and harmonized to create one final object with 73 unique communities/cell states at the most granular level.

#### Annotation of Neutrophil subsets

To further analyze the neutrophil compartment, we harmonized and clustered all cells labeled as neutrophils as above. We then performed marker gene identification for each of the clusters and manually merged clusters until we could identify marker genes specific to each major subset of cells. Neutrophil subsets were determined by looking at the top 10 marker genes for each subgroup and identifying known neutrophil-associated genes.

#### Differential Abundance Analysis

We performed differential abundance analysis to compare the number of cells identified between uninflamed and inflamed conditions in our labeled single-cell data subsets. Differential abundance calculates the total number of cells per condition at the per-sample level and tests the null hypothesis that the mean abundance between uninflamed and inflamed samples equals zero. We tested this hypothesis using edgeR’s gene-wise negative binomial generalized linear model with a quasi-likelihood test (60). Briefly, we first converted our Seurat object to a single-cell experiment object and then calculated the abundances of each cell type using base R’s table function. We did not filter out cell types with low expression counts since this would lose specific cell subsets with lower numbers of cells. We then created a model matrix design using uninflamed vs. inflamed as our covariate in addition to patient samples. We then estimated common, trended, and tagwise negative binomial dispersions by weighted likelihood empirical bayes using the estimateDisp function. Next, we scaled our raw library sizes using edgeR’s calcNormFactors to account for differences in library sample sizes. We then fit our data to a generalized linear model with a negative binomial distribution using the edgeRs glmQLFit function. Lastly, we performed a negative binomial test using edgeRs glmQLFTest function on our fitted data and considered cell types as differentially abundant if they had a false discovery rate value of less than 0.05.

#### Derivation and application of cell-specific gene modules

Cell-specific gene modules were derived from the scRNA-seq data by first identifying marker genes for each cell population using the Wilcoxauc function from the presto package (61). For each cell population, an initial gene module was created by combining the top 20 marker genes by logFC on cells of interest with the top 20 marker genes by the percentage of cells in the population of interest expressing the gene filtered for markers expressed by at most 2 percent of the other cells. This initial list was further manually refined by removing any non-specific marker genes.

Cell-specific signature scores for each sample were calculated by taking the average of logCPM for the corresponding gene module. Analysis of the association of longitudinal changes with MCS remission or treatment was performed using the voom-limma framework, with the subject introduced as a blocking factor and adjusted for patient-reported sex.

#### ROC analysis

A logistic regression model was developed to evaluate the association between MCS remission at induction with the longitudinal log2 fold change for each neutrophil state, adjusted for patient reported sex. Similarly, another logistic regression model was built to assess the association between MCS remission at induction with the baseline levels of each neutrophil state, adjusted for patient-reported sex. The ROC curves were generated by comparing the observed and the predicted MCS remission status based on each regression model.

#### Trajectory analysis

We performed trajectory analysis using Monocle 3 (62–64) to identify differences in neutrophil subset states. We first created a Seurat object that contained only neutrophils and excluded low RNA neutrophils, which we reharmonized and reclustered to generate new UMAP dimension reduction values. We then converted our Seurat object to the native Monocle 3 data format and estimated size factors. Next, we clustered the cells using the Leiden clustering algorithm with a *k* value 30 using Monocle3’s cluster_cells function. Additionally, we identified principal graphs from the reduced dimension space using reversed graph embedding with the learn_graph function. To identify root nodes for the origin of our pseudotime trajectories in an unbiased fashion, we used Monocle3’s get_earlierst_principal_node function when ordering cells. This assigned a pseudotime trajectory to all cells within our neutrophil lineage, which we then used for downstream analysis.

To visualize the expression of genes across pseudotime, we first subset our dataset to contain only the genes of interest, including CXCR2 and CXCR4. We then generated a plot showing the expression of genes as a function of pseudotime using the plot_genes_in_pseudotime function within Monocle 3, filtering out for a minimum expression value of 0.5 for each of these genes to remove lowly expressing cells. We used the default formula, which uses a natural cubic spline with a degree of freedom of 3 using the *ns* function.

To generate a heatmap that shows gene expression across pseudotime for multiple genes, we tested all genes for differential expression using the graph_test function on the principal graph using our UMAP representation. This function tests for differential expression based on the low dimensional embedding and principal graph of the scRNA-seq data by calculating p-values, q-values, and Moran’s I values for each gene. We then set a threshold for both a q value < 0.001 and a Moran’s I value > 0.05 to loosely filter for genes that significantly varied over pseudotime. We fit a spline with 3 degrees of freedom using the smooth.spline function from the stats package to the log2 count matrix of genes passing these cutoffs. We then converted this matrix to Z-score values and generated bins for cells across pseudotime containing 100 cells per bin. The binned matrix was used to create a pseudotime expression heatmap. To summarize genes that varied across pseudotime, we reported both the maximum expression of each gene and the bin in which we observed maximum expression.

#### Cell-cell interactions

Cell-cell interaction analysis was performed using CellChat (65). A Seurat object with all cell types except neutrophil-low RNA was created. We then created cellChat objects for inflamed and uninflamed tissue and a merged object of inflamed vs. uninflamed tissue for differential analysis. IL1B: IL1R1 interaction was not within the cellChatDB database and was manually added based on evidence from (66). The minimum number of cells to create the communication network for a cell population was set to 10. We did not create interactions of Adventitial fibroblast in the inflamed tissue due to low cell numbers (n=8). Analyses were run using default parameters.

### Data availability

Data will be made available to qualified researchers on acceptance of the manuscript for publication.

---

**Figure S1.**
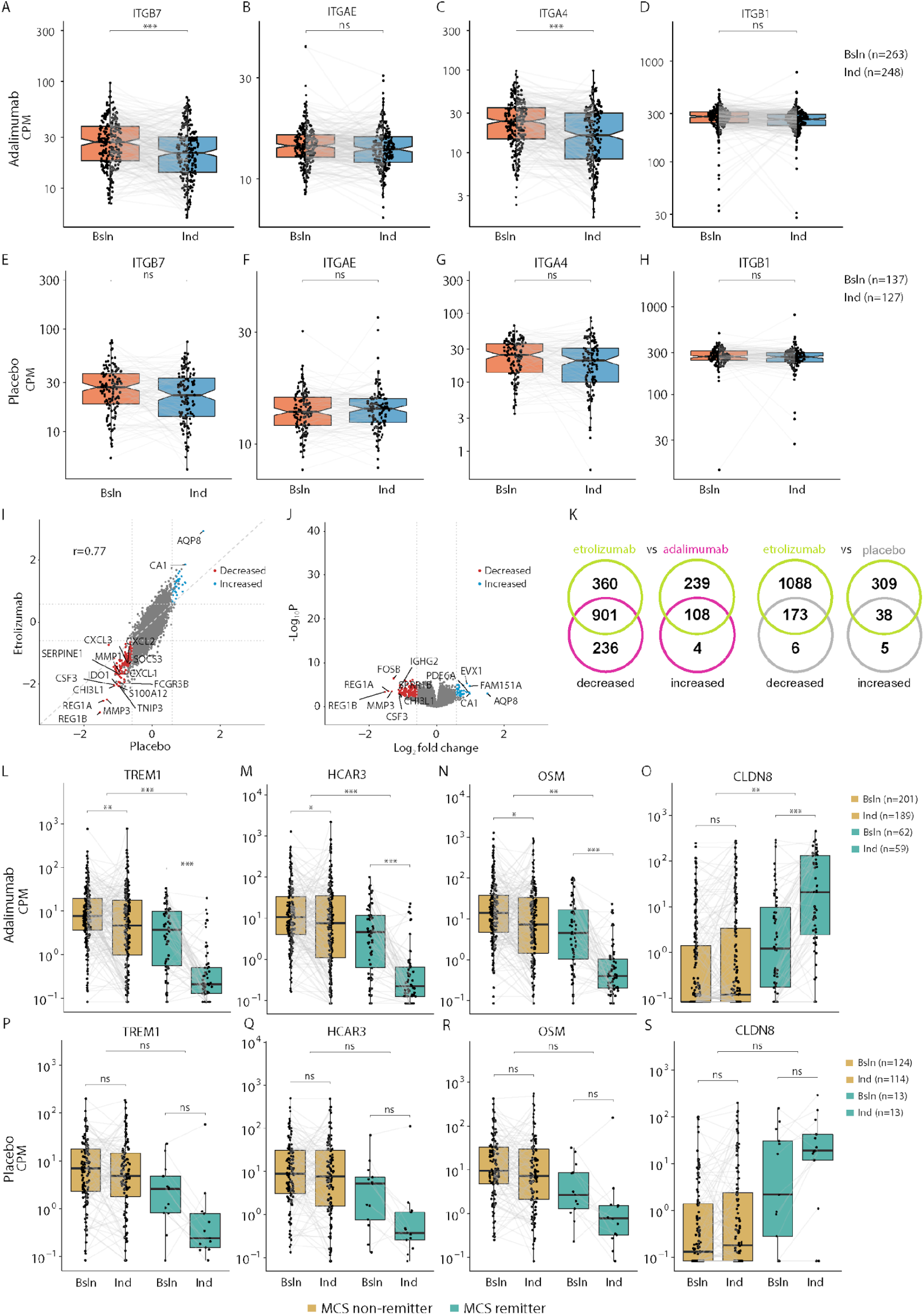
Transcriptomic analysis of etrolizumab, adalimumab, and placebo treatment in colonic biopsies. **A.-D.** Expression of selected integrins before (baseline) and after (induction; week 10) adalimumab treatment. Each point represents an individual patient, and pre-and post-treatment trajectories for each patient are shown (Gray bar). Expression values are in normalized counts per million (CPM). Box plots represent the upper and lower quartiles, the middle line represents the median, notches show 1.58x the interquartile range (IQR) divided by the square root of the number of samples measured, and the whiskers extend to the most extreme point no more than 1.5x outside of the interquartile ranges. **E.-H.** Expression of selected integrins before (baseline) and after (induction; week 10) placebo treatment. **I.** Comparison of log2 fold changes between placebo (x-axis) and etrolizumab (y-axis). Each point represents a gene, colored blue for genes significantly up-regulated after treatment (fold change > 1.5x at an FDR of 0.05) and colored red for genes significantly down-regulated after treatment (fold change < −1.5x at an FDR of 0.05). **J.** Volcano plots showing log2 fold change on the x-axis and -log10 p-value on the y-axis for changes in gene expression after placebo treatment for 10 weeks. Each point represents a gene, colored as in **I.** above. **K.** Venn diagrams comparing the number of genes significantly down-regulated or up-regulated after treatment with etrolizumab or adalimumab for 10 weeks and the number of genes significantly down-regulated or up-regulated after treatment with etrolizumab or placebo for 10 weeks. **L.-O.** Expression of selected neutrophil-associated or epithelium-associated genes before (baseline) or after (induction; week 10) adalimumab treatment. Each point represents a sample collected from a patient at the respective time point, and patient measurements are joined by a line. Boxes are colored by remission status of the patient at week 10. Boxes represent the upper and lower quartile, the middle line represents the median, and the whiskers extend to the most extreme point no more than 1.5x the interquartile range from the box boundaries. For all panels, stars indicate Benjamini-Hochberg corrected p-values: ns: p > 0.01; *: p < 0.01; ** p < 0.001; *** p < 0.0001. **P.-S.** Expression of selected neutrophil-associated or epithelium-associated genes before (baseline) or after (induction; week 10) placebo treatment.

**Figure S2.**
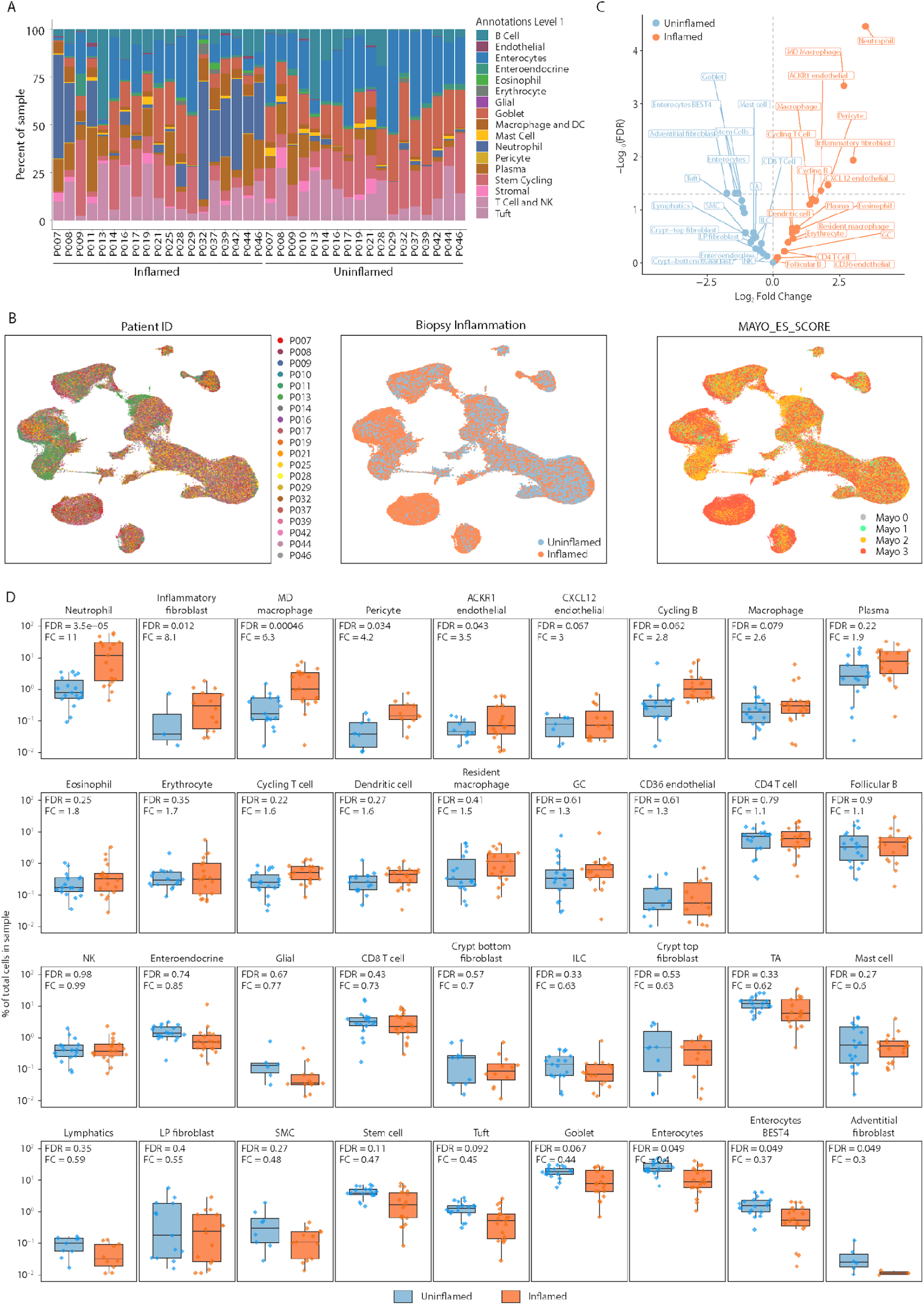
Neutrophils are elevated in inflamed and more severe patient samples. **A. Patient** cell type percentage distribution across inflamed and uninflamed samples at the broadest level of annotation. Each bar represents a sample of colonic tissue from a patient, and colors indicate the percent of cell type within stacked barplot n = 19 inflamed and 18 uninflamed. **B.** Summary statistics for differential abundance analysis at Annotation level 2 are shown. Dotplot depicting log2 fold change (x-axis) and -log10(FDR) (y-axis) for each cell type at annotation level 2 across all patients. Dots in blue are elevated in uninflamed samples, and dots in orange are elevated in inflamed samples. All dots above the dashed line are significantly elevated with an FDR value of < 0.05. **C.** Uniform manifold approximation plots (UMAPs) showing Patient ID, Biopsy inflammation status, and Mayo Endoscopy subscore across all patients. **D.** Box plots ordered by FDR, depicting differential abundance analysis at the patient level for all cell types at an intermediate level of granularity, split by inflammation status. The y-axis represents the percentage of each cell type out of the total cells in the sample. Each dot represents a sample from a patient. Blue boxes represent uninflamed samples, and orange boxes represent inflamed samples. FDR and fold change are displayed in the upper left corner of each plot and were calculated separately (see Methods).

**Figure S3.**
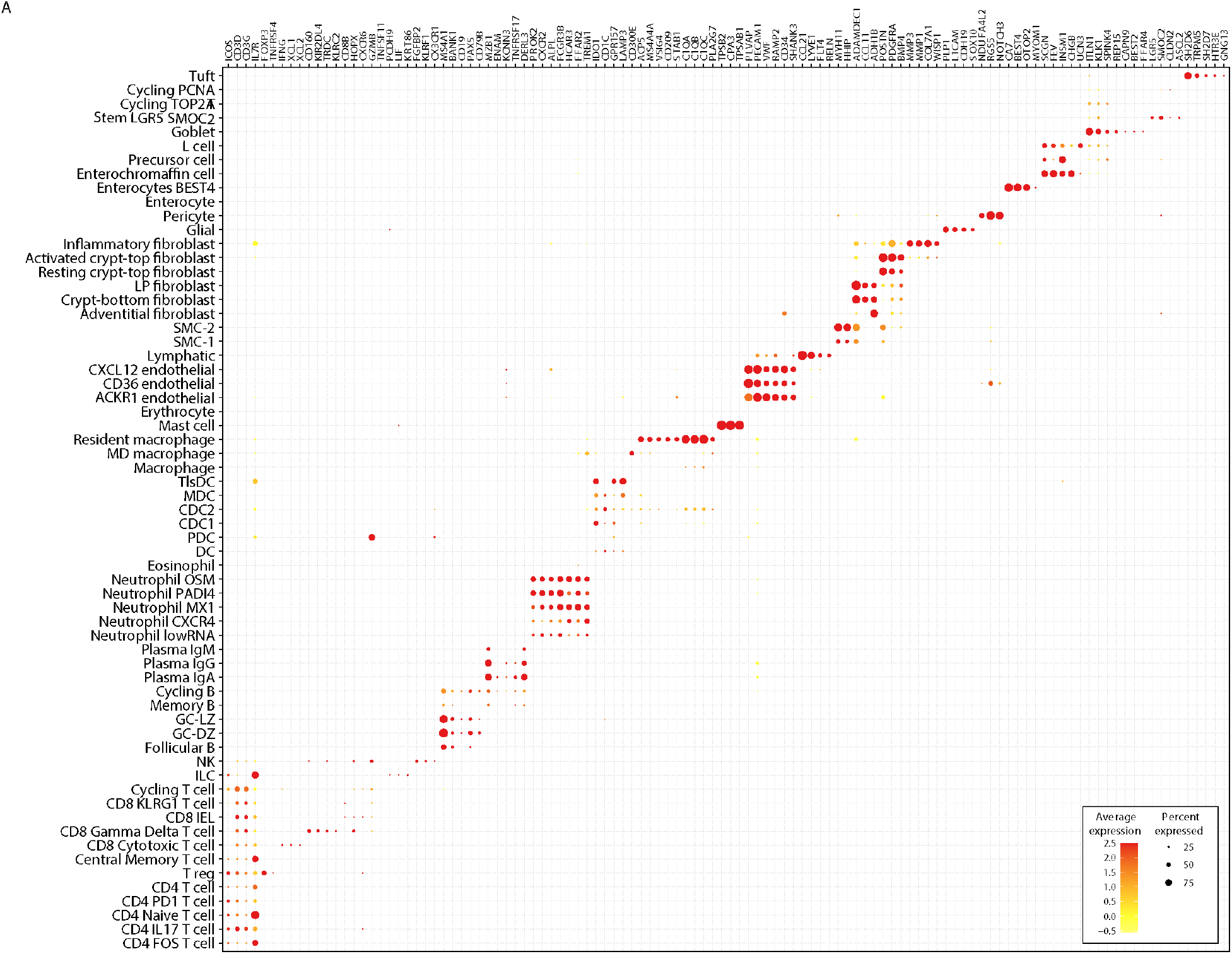
Single-cell gene signatures derived to deconvolve bulk transcriptomics data. **A.** Single-cell-derived cell-specific gene modules applied to bulk transcriptomics data are shown. Each point represents the expression of a gene in a cell population. The size of the point indicates the proportion of cells expressing the gene, and the color indicates the centered and scaled expression level.

**Figure S4.**
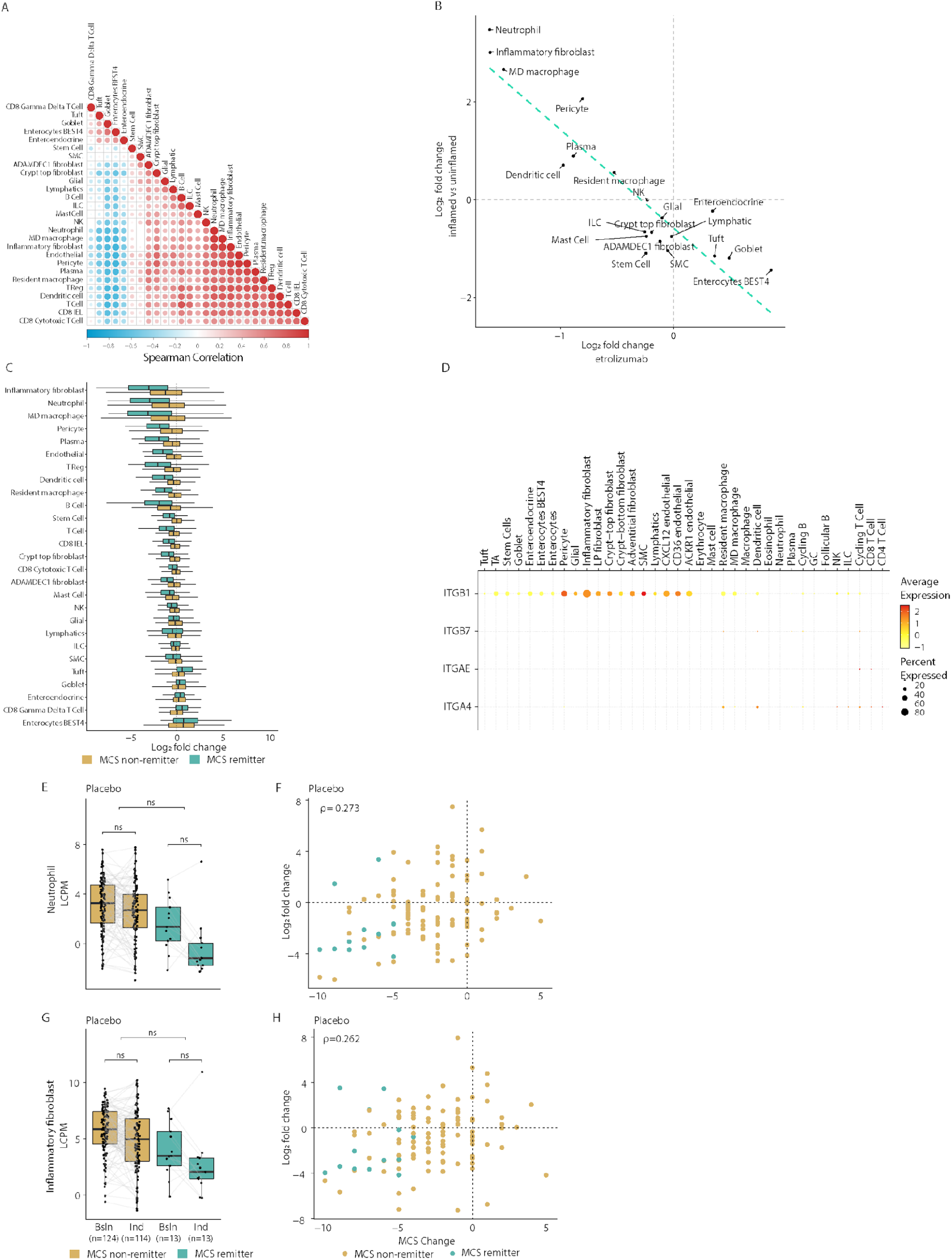
Cellular specificity of selected integrins in UC tissue and correlation between and changes in gene modules in adalimumab and placebo-treated patients. **A.** Heatmap of Spearman correlation between signature scores for individual samples. **B.** Effect of etrolizumab on individual cell type expression signatures (x-axis) compared with differential abundance of the cell type in inflamed vs uninflamed biopsies (y-axis). Each point represents an individual cell population. **C.** Effect of treatment on genes specific for individual cell populations in remitters or non-remitters after adalimumab treatment. Boxes represent the log2 fold change between baseline and week 10 for either non-remitters or remitters. Cell populations are sorted according to the order in Figure 3B. **D.** Each point represents the expression of an integrin gene on a cell population. The size of the point indicates the proportion of cells expressing the gene, and the color indicates the centered and scaled expression level. **E.** Expression of neutrophil signature score before and after treatment with placebo. Each point represents a patient sample before (baseline) or after (induction, week 10) treatment. Samples collected from the same patient are linked by a line. **F.** Expression of inflammatory fibroblast signature score before and after treatment with placebo. Each point represents a patient sample before (baseline) or after (induction, week 10) treatment. Samples collected from the same patient are linked by a line. **G.** Comparison of log2 fold change in neutrophil gene expression signature (y-axis) with change in Mayo Clinic score (x-axis) after treatment with placebo. Each point represents an individual patient, colored by remission status at week 10. **H.** Comparison of log2 fold change in inflammatory fibroblast gene expression signature (y-axis) with change in Mayo Clinic score (x-axis) after treatment with placebo. Each point represents an individual patient, colored as in D.

**Figure S5.**
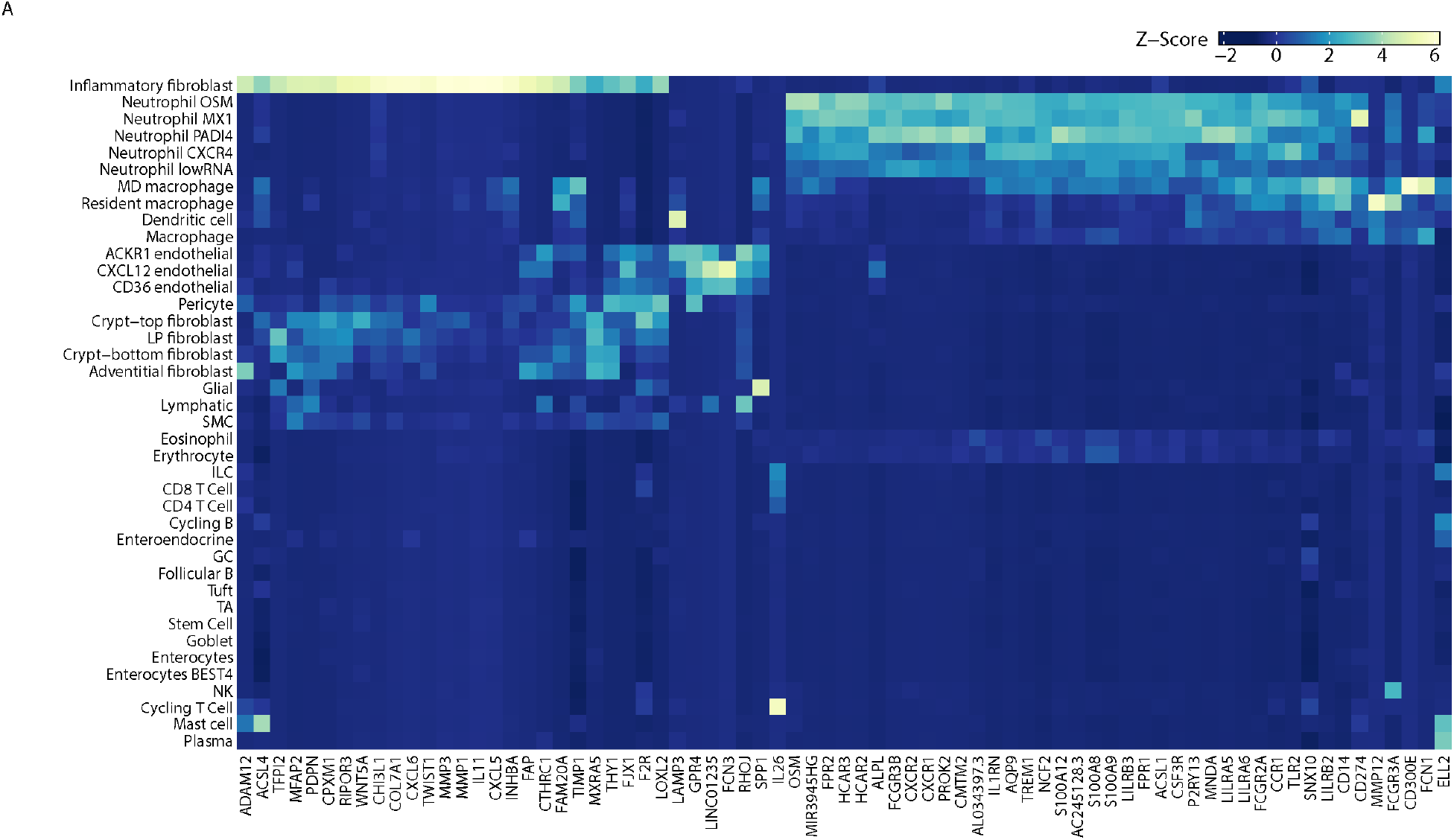
Anti-TNF resistance modules are expressed by inflammatory fibroblast and neutrophil subsets. **A.** Heatmap showing the Z-Score of average expression of TNF resistance module genes M4 and M5 from Friedrich et al. (12) at level 2 annotations.

**Figure S6.**
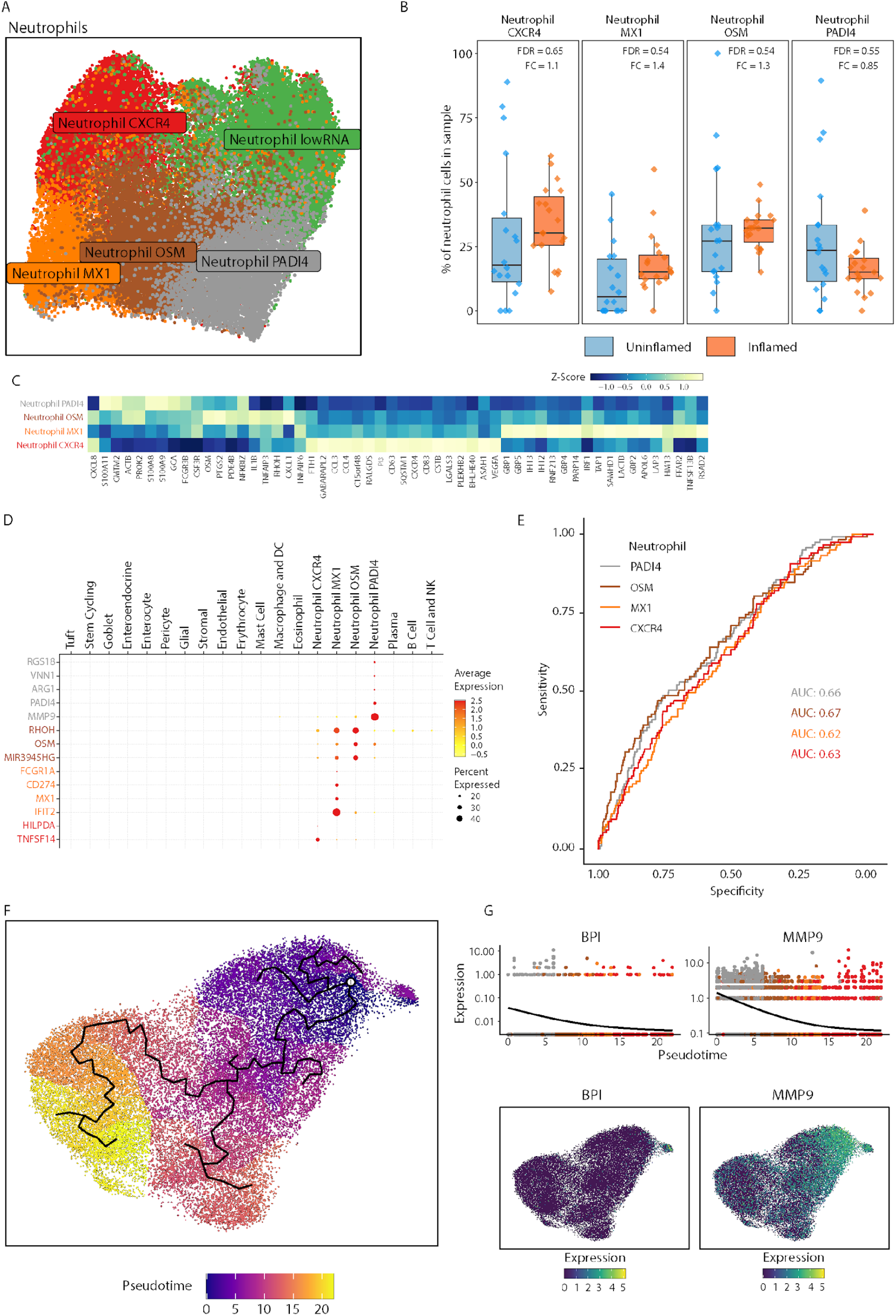
Heterogeneity of neutrophils in UC tissue biopsies. **A.** UMAP projection of neutrophil subsets, including low RNA neutrophils. Each point represents a cell, colored by the respective subset it was classified as. **B.** Differential abundance of neutrophils across inflamed and uninflamed tissue. Each point represents a sample. The y-axis shows the fraction of each neutrophil subset relative to the total number of neutrophils in that sample. Fold changes and FDR values were determined as described in the methods. **C.** Heatmap showing the Z-Score of average expression of genes used to identify neutrophil subsets from Garrido-Trigo et al. (18) mapped onto our neutrophil subsets. Rows are ordered from the earliest subset to the latest subset in pseudotime, and columns are hierarchically clustered using Euclidean distance. **D.** Single cell-derived gene modules specific to neutrophil subsets are shown across all cell populations. Each point represents the expression of a gene in a cell population. The size of the point indicates the proportion of cells expressing the gene, and the color indicates the centered and scaled expression level. **E.** ROC curves for the association of baseline levels of neutrophil subsets with remission. **F.** UMAP projection of neutrophil subsets colored by the inferred pseudotime. The inferred trajectory is shown in black lines. The white circle indicates the starting point for the trajectory inference **G.** Expression of BPI and HP along the neutrophil pseudotime axis. Each dot represents a cell with black lines indicating the expression of each gene along pseudotime. The UMAPs below show the expression of BPI and MMP9 in neutrophils.

**Figure S7.**
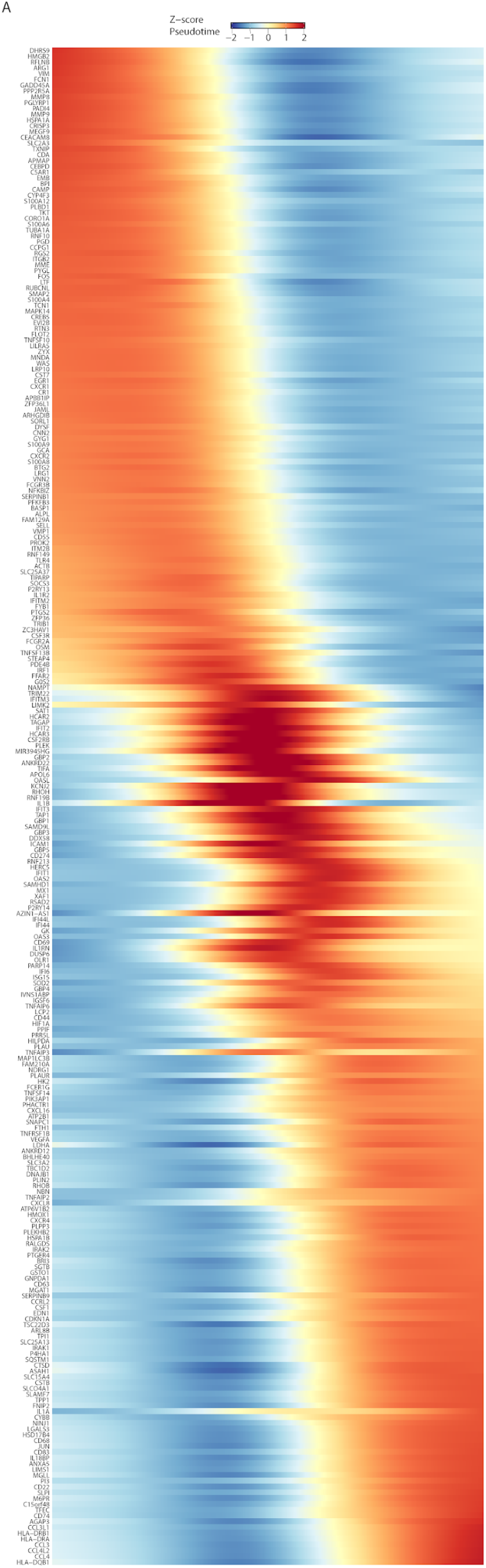
Gene expression across pseudotime for neutrophil subsets. **A.** Heatmap showing the Z-Score of log normalized gene expression values for genes with a q-value of < 0.001 and a Moran’s I value > 0.05 after testing differential gene expression across pseudotime. Genes are ordered across pseudotime. Pseudotime consists of bins, each containing 100 cells.

**Figure S8.**
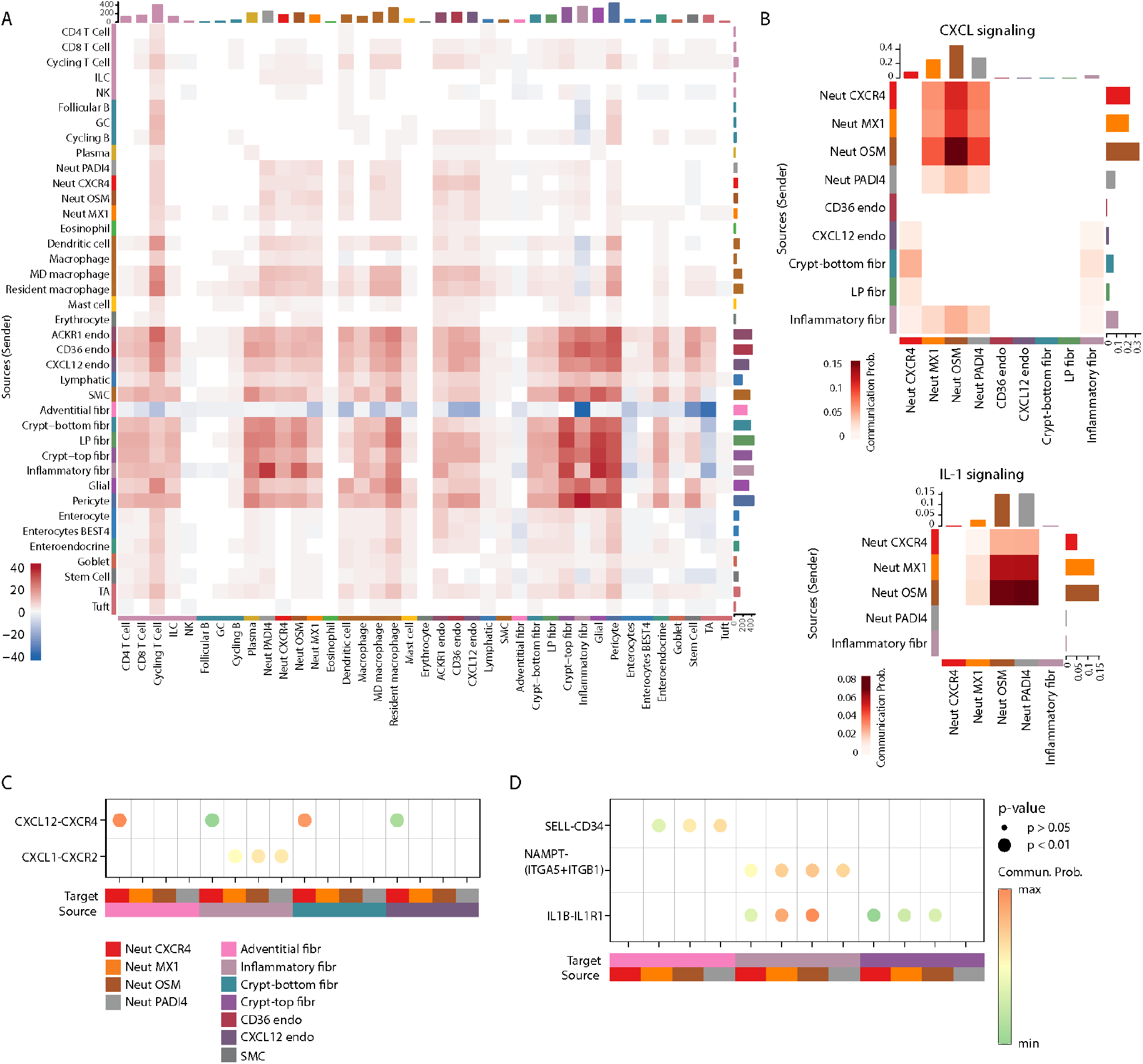
Neutrophil interactions in the UC uninflamed tissue. **A.** Differential number of inferred interactions between inflamed and uninflamed tissue based on cellChat. Adventitial fibroblast cells in inflamed tissue were excluded from cell-cell interaction analysis due to the low number of cells identified (8 cells). **B.** Heatmap of signaling pathways CXCL and IL1 showing significant communications between neutrophil subpopulations and fibroblasts and endothelial. We observe fewer communication probabilities in the uninflamed tissue than in the inflamed tissue. The color in the heatmap is proportional to the overall communication probability between each cell type. **C.** Selected significant ligand-receptors as incoming signaling interactions to neutrophil subpopulations from fibroblasts and endothelial subsets in the uninflamed tissue. **D.** Significant ligand-receptors were selected as outgoing signaling interactions between neutrophil subpopulations, fibroblasts, and endothelial subsets in the uninflamed tissue.

## References

1. S. C. Ng, H. Y. Shi, N. Hamidi, F. E. Underwood, W. Tang, E. I. Benchimol, R. Panaccione, S. Ghosh, J. C. Y. Wu, F. K. L. Chan, J. J. Y. Sung, G. G. Kaplan, Worldwide incidence and prevalence of inflammatory bowel disease in the 21st century: a systematic review of population-based studies. Lancet, 390, 2769–2778 (2018).

2. B. Click, C. R. Rivers, I. E. Koutroubakis, D. Babichenko, A. M. Anderson, J. G. Hashash, M. A. Dunn, M. Schwartz, J. Swoger, L. Baidoo, A. Barrie, M. Regueiro, D. G. Binion, Demographic and Clinical Predictors of High Healthcare Use in Patients with Inflammatory Bowel Disease. Inflamm.Bowel Dis. 22, 1442–1449 (2016).

3. R. Ungaro, S. Mehandru, P. B. Allen, L. Peyrin-Biroulet, J.-F. Colombel, Ulcerative colitis. Lancet 389, 1756–1770 (2017).

4. M. F. Neurath, Current and emerging therapeutic targets for IBD. Nat. Rev. Gastroenterol. Hepatol. 14, 269–278 (2017).

5. A. C. Moss, Optimizing the use of biological therapy in patients with inflammatory bowel disease. Gastroenterol. Rep. 3, 63–68 (2015).

6. S. Vermeire, S. O’Byrne, M. Keir, M. Williams, T. T. Lu, J. C. Mansfield, C. A. Lamb, B. G. Feagan, J. Panes, A. Salas, D. C. Baumgart, S. Schreiber, I. Dotan, W. J. Sandborn, G. W. Tew, D. Luca, M. T. Tang, L. Diehl, J. Eastham-Anderson, G. D. Hertogh, C. Perrier, J. G. Egen, J. A. Kirby, G. V. Assche, P. Rutgeerts, S. O. ’byrne, M. Keir, M. Williams, T. T. Lu, J. C. Mansfi, C. A. Lamb, B. G. Feagan, J. Panes, A. Salas, D. C. Baumgart, S. Schreiber, I. Dotan, W. J. Sandborn, G. W. Tew, D. Luca, M. T. Tang, L. Diehl, J. R. Eastham-Anderson, G. D. Hertogh, C. Perrier, J. G. Egen, J. A. Kirby, G. V. Assche, P. Rutgeerts, Etrolizumab as induction therapy for ulcerative colitis: A randomised, controlled, phase 2 trial. Lancet 384, 309–318 (2014).

7. B. Dai, J. A. Hackney, R. Ichikawa, A. Nguyen, J. Elstrott, L. D. Orozco, K.-H. Sun, Z. Modrusan, A. Gogineni, A. Scherl, J. Gubatan, A. Habtezion, M. Deswal, M. Somsouk, W. A. Faubion, A. Chai, Z. Sharafali, A. Hassanali, Y. S. Oh, S. Tole, J. McBride, M. E. Keir, T. Yi, Dual targeting of lymphocyte homing and retention through α4β7 and αEβ7 inhibition in inflammatory bowel disease. Cell Rep Med. 2, 100381 (2021).

8. M. E. Keir, F. Fuh, R. Ichikawa, M. Acres, J. A. Hackney, G. Hulme, C. D. Carey, J. Palmer, C. J. Jones, A. K. Long, J. Jiang, S. Klabunde, J. C. Mansfield, C. M. Looney, W. A. Faubion, A. Filby, J. A. Kirby, J. McBride, C. A. Lamb, Regulation and Role of αE Integrin and Gut Homing Integrins in Migration and Retention of Intestinal Lymphocytes during Inflammatory Bowel Disease. J Immunol. 207 (9): 2245–2254(2021)

9. D. T. Rubin, I. Dotan, A. DuVall, Y. Bouhnik, G. Radford-Smith, P. D. R. Higgins, D. S. Mishkin, P. Arrisi, A. Scalori, Y. S. Oh, S. Tole, A. Chai, K. Chamberlain-James, S. Lacey, J. McBride, J. Panés, A. Rustem, A. B. Norasiah, A. Humberto, A. Diego, A. Hale, A. Evangelos, A. Olga, A. Bagdadi, A. Andres, A. Ashwin, A. Jane, A. Tomasz, A. Nathan, A. Ozlen, B. Mauro, A. Jozef, B. George, B. Marko, B. Andrey, B. N. Guerino, B. Metin, B. Curtis, B. Stefan, B. William, B. Fatih, B. Sudhir, B. Andrzej, B. Leonid, B. Bahri, B. Pavol, B. Verle, B. Julia, B. Vladimir, B.-P. Francisco, B. Yoram, B. James, B. Tetiana, B. Igor, B. Ivan, C. Jonathon, C. A. Azlida, C. Tatiana, C. Michael, C. Ivan, C. Dimitrios, C. P. Shan, C. Galina, C. Andrew, C. Robert, C. Mirjana, D. Ulku, D. W. Cezary, D. Olena, D. Henry, D. Elena, D. Jelena, D. J. Nik, D. Julia, D. Oleg, D. Tomas, D. Piotr, D. Gerald, D. G. Pedro, D. G. Aaron, D. Mikhail, E. Craig, E. Yusuf, F. Galyna, F. Oleksandr, F. Olga, F. Miroslav, F. Roland, F. Jorge, F. M. Lorena, F. Lucky, F. Bradley, F. Keith, F. Sergio, G.-W. Beata, G. P. F. Leonel, G. E. Daniel, G. Nataliia, G. Oleksandr, G. Maciej, G. Can, G. Glenn, G. Milos, G. Vladimir, G. R. Rogelio, H. Stephen, H. John, H. Marek, H. Xavier, H. Peter, H. Robert, H. David, H. Peter, H. Raouf, H. I. Normiha, H. Tibor, H. Richard, H. Gerald, H. John, H. Frantisek, H. Ihor, H. Irena, H. Sadettin, I. V. L. Alberto, I. Ikechukwu, I. Stephen, I. Vladimir, I. James, J. Rajesh, J.-K. Zofia, K. Victor, K. John, K. Tarkan, K. Marek, K. Irina, K. Seymour, K. Barry, K. Edita, K. Irina, K. Sunil, K. Jaroslaw, K. Anzela, K. Dariusz, K. Volodymyr, K. Slavko, K.-S. Malgorzata, K. Natalya, K. Bartosz, K. Lyubomir, K. Iskren, K. Georgios, K. Ioannis, K. Richard, K. Ian, K. Miodrag, K. Zeljko, K. Mikolaj, K. Grazyna, K. Karin, K. Limas, L. Mark, L. R. Tatjana, L. Rupert, L. W. Keung, L. Henry, L. M. K. Kong, L. B. C. B. Lúcia, L. Maria, L. Tetiana, L. P. M. Helena, L. John, L. Kresimir, L. Milan, L. Yurii, M. Finlay, M. Anu, M. Igor, M. Arkadiusz, M. Gerasimos, M. Benno, M. Ivanka, M. Inna, M. Mario, M. Srdjan, M. V. J. Ricardo, M. Felipe, M. Konstantinos, M. Brent, M. Gregory, M. G. L. Alonso, M. Salvatore, M. Yuriy, M. Reme, N. Aleksandar, N. Viacheslav, O. Andrey, O. Oleksandr, O. S. S. Genoile, O. Maria, P. Vladimir, P. Dimitar, P. Mariana, P. Sasa, P. Plamen, P. Asen, P. Plamen, P. Michaela, P. Raymond, P. C. Sergio, P. Igor, P. Ludmyla, P. Mykhailo, P. C. A. Teresa, P. Aldis, P. Jiri, P. Volodymyr, R. Istvan, R.-S. Graham, R. A. R. Affendi, R. C. Daniel, R. J. Odery, R. David, R. Andrey, R. Jaroslaw, R. Amir, R. Viktoriia, R. Iaroslava, R. Xavier, R. Grigory, R. R. C. Arturo, R. Jerzy, R. David, R. Maciej, R. Jacek, S. Oleg, S. Halil, S. Rosemi, S. Douglas, S. S. Antonio, S. Robert, S. Michael, S. Michael, S. John, S. Shahriar, S. Ahmad, S. Marina, S. Natalia, S. Oksana, S. Alex, S. Irina, S. Vladimir, S. Vladislav, S. Giedrius, S. Igor, S. Zbigniew, S. Jan, S. Mahmood, S. Najm, S. Konstantinos, S. Zoia, S. Mykola, S.-M. Krystyna, S. Jonathas, S. Simeon, S. Girgina, S. Keith, S. Lindsey, T. Dimitar, T. Jaak, T. Ludmila, T. Hugo, T. Dino, T. Elias, T. Konstantin, T. H. Poh, T. Lena, T. Carlton, T. Irina, T. Tsveta, T. Oleksandr, T. Ivars, T. Ratko, T. Vasiliy, T. Zsolt, U. Belkis, U.-G. Alma, V. John, V. Ekaterina, V. Eduardo, V. Galina, V. Sergiy, V. Byron, V. Francisco, V. Vadym, V. Borislav, V. Miroslava, V. Petr, W. Ian, W. Marek, W. William, W. John, W.-K. Anna, W. Nathaniel, W. Pawel, W.-S. Barbara, Y. Bruce, Y. Alexey, Y. Ziad, Y. S. Lígia, Y. Ilhami, Z. Jan, Z. Cyrla, Z. P. Natasa, Z. Vyacheslav, Z. Maryna, Z. Maciej, Etrolizumab versus adalimumab or placebo as induction therapy for moderately to severely active ulcerative colitis (HIBISCUS): two phase 3 randomised, controlled trials. Lancet Gastroenterol Hepatol. 7, 17–27 (2022).

10. I. Arijs, K. Li, G. Toedter, R. Quintens, L. V. Lommel, K. V. Steen, P. Leemans, G. D. Hertogh, K. Lemaire, M. Ferrante, F. Schnitzler, L. Thorrez, K. Ma, X.-Y. R. Song, C. Marano, G. V. Assche, S. Vermeire, K. Geboes, F. Schuit, F. Baribaud, P. Rutgeerts, Mucosal gene signatures to predict response to infliximab in patients with ulcerative colitis. Gut 58, 1612–1619 (2009).

11. C. S. Smillie, M. Biton, J. Ordovas-Montanes, K. M. Sullivan, G. Burgin, D. B. Graham, R. H. Herbst, N. Rogel, M. Slyper, J. Waldman, M. Sud, E. Andrews, G. Velonias, A. L. Haber, K. Jagadeesh, S. Vickovic, J. Yao, C. Stevens, D. Dionne, L. T. Nguyen, A.-C. Villani, M. Hofree, E. A. Creasey, H. Huang, O. Rozenblatt-Rosen, J. J. Garber, H. Khalili, A. N. Desch, M. J. Daly, A. N. Ananthakrishnan, A. K. Shalek, R. J. Xavier, A. Regev, Intra- and Inter-cellular Rewiring of the Human Colon during Ulcerative Colitis. Cell 178, 714–730.e22 (2019).

12. M. Friedrich, M. Pohin, M. A. Jackson, I. Korsunsky, S. J. Bullers, K. Rue-Albrecht, Z. Christoforidou, D. Sathananthan, T. Thomas, R. Ravindran, R. Tandon, R. S. Peres, H. Sharpe, K. Wei, G. F. M. Watts, E. H. Mann, A. Geremia, M. Attar, F. Barone, M. Brenner, C. D. Buckley, M. Coles, A. P. Frei, K. G. Lassen, F. M. Powrie, S. McCuaig, L. Thomas, E. Collantes, H. H. Uhlig, S. N. Sansom, A. Easton, S. Raychaudhuri, S. P. Travis, F. M. Powrie, IL-1-driven stromal–neutrophil interactions define a subset of patients with inflammatory bowel disease that does not respond to therapies. Nat. Med. 27, 1970–1981(2021).

13. D. Jha, Z. Al-Taie, A. Krek, S. T. Eshghi, A. Fantou, T. Laurent, M. Tankelevich, X. Cao, H. Meringer, A. E. Livanos, M. Tokuyama, F. Cossarini, A. Bourreille, R. Josien, R. Hou, P. Canales-Herrerias, R. C. Ungaro, M. Kayal, J. Marion, A. D. Polydorides, H. M. Ko, D. D’souza, R. Merand, S. Kim-Schulze, J. A. Hackney, A. Nguyen, J. M. McBride, G.-C. Yuan, J. F. Colombel, J. C. Martin, C. Argmann, M. Suárez-Fariñas, F. Petralia, S. Mehandru, Myeloid cell influx into the colonic epithelium is associated with disease severity and non-response to anti-Tumor Necrosis Factor Therapy in patients with Ulcerative Colitis. [Preprint]. 2023 Jun 5:2023.06.02.542863. doi: 10.1101/2023.06.02.542863.

14. J. C. Martin, C. Chang, G. Boschetti, R. Ungaro, M. Giri, J. A. Grout, K. Gettler, L. Chuang, S. Nayar, A. J. Greenstein, M. Dubinsky, L. Walker, A. Leader, J. S. Fine, C. E. Whitehurst, M. L. Mbow, S. Kugathasan, L. A. Denson, J. S. Hyams, J. R. Friedman, P. T. Desai, H. M. Ko, I. Laface, G. Akturk, E. E. Schadt, H. Salmon, S. Gnjatic, A. H. Rahman, M. Merad, J. H. Cho, E. Kenigsberg, Single-Cell Analysis of Crohn’s Disease Lesions Identifies a Pathogenic Cellular Module Associated with Resistance to Anti-TNF Therapy. Cell 178, 1493–1508.e20 (2018).

15. N. R. West, A. N. Hegazy, B. M. J. Owens, S. J. Bullers, B. Linggi, S. Buonocore, M. Coccia, D. Görtz, S. This, K. Stockenhuber, J. Pott, M. Friedrich, G. Ryzhakov, F. Baribaud, C. Brodmerkel, C. Cieluch, N. Rahman, G. Müller-Newen, R. J. Owens, A. A. Kühl, K. J. Maloy, S. E. Plevy, S. Keshav, S. P. L. Travis, F. Powrie, C. Arancibia, A. Bailey, E. Barnes, B. Bird-Lieberman, O. Brain, B. Braden, J. Collier, J. East, L. Howarth, S. Keshav, P. Klenerman, S. Leedham, R. Palmer, F. Powrie, A. Rodrigues, A. Simmons, P. Sullivan, H. Uhlig, Oncostatin M drives intestinal inflammation and predicts response to tumor necrosis factor–neutralizing therapy in patients with inflammatory bowel disease. Nat. Med. 23, 579–589 (2017).

16. D. Aschenbrenner, M. Quaranta, S. Banerjee, N. Ilott, J. Jansen, B. Steere, Y.-H. Chen, S. Ho, K. Cox, C. V. Arancibia-Cárcamo, M. Coles, E. Gaffney, S. P. Travis, L. Denson, S. Kugathasan, J. Schmitz, F. Powrie, S. N. Sansom, H. H. Uhlig, Deconvolution of monocyte responses in inflammatory bowel disease reveals an IL-1 cytokine network that regulates IL-23 in genetic and acquired IL-10 resistance. Gut 70, 1023–1036 (2021).

17. Y. Haberman, R. Karns, P. J. Dexheimer, M. Schirmer, J. Somekh, I. Jurickova, T. Braun, E. Novak, L. Bauman, M. H. Collins, A. Mo, M. J. Rosen, E. Bonkowski, N. Gotman, A. Marquis, M. Nistel, P. A. Rufo, S. S. Baker, C. G. Sauer, J. Markowitz, M. D. Pfefferkorn, J. R. Rosh, B. M. Boyle, D. R. Mack, R. N. Baldassano, S. Shah, N. S. Leleiko, M. B. Heyman, A. M. Grifiths, A. S. Patel, J. D. Noe, B. J. Aronow, S. Kugathasan, T. D. Walters, G. Gibson, S. D. Thomas, K. Mollen, S. Shen-Orr, C. Huttenhower, R. J. Xavier, J. S. Hyams, L. A. Denson, Ulcerative colitis mucosal transcriptomes reveal mitochondriopathy and personalized mechanisms underlying disease severity and treatment response. Nat. Commun. 10, 38 (2019).

18. A. Garrido-Trigo, A. M. Corraliza, M. Veny, I. Dotti, E. Melón-Ardanaz, A. Rill, H. L. Crowell, Á. Corbí, V. Gudiño, M. Esteller, I. Álvarez-Teubel, D. Aguilar, M. C. Masamunt, E. Killingbeck, Y. Kim, M. Leon, S. Visvanathan, D. Marchese, G. Caratù, A. Martin-Cardona, M. Esteve, J. Panés, E. Ricart, E. Mereu, H. Heyn, A. Salas, Macrophage and neutrophil heterogeneity at single-cell spatial resolution in human inflammatory bowel disease. Nat. Commun. 14, 4506 (2023).

19. S. Singh, A. N. Ananthakrishnan, N. H. Nguyen, B. L. Cohen, F. S. Velayos, J. M. Weiss, S. Sultan, S. M. Siddique, J. Adler, K. A. Chachu, A. C. G. Committee, AGA Clinical Practice Guideline on the Role of Biomarkers for the Management of Ulcerative Colitis. Gastroenterology 164, 344–372 (2023).

20. J. Wang, M. Macoritto, H. Guay, J. W. Davis, M. C. Levesque, X. Cao, The Clinical Response of Upadacitinib and Risankizumab Is Associated With Reduced Inflammatory Bowel Disease Anti-TNF-α Inadequate Response Mechanisms. Inflamm. Bowel Dis. 29, 771–782 (2022).

21. H. M. Penrose, R. Iftikhar, M. E. Collins, E. Toraih, E. Ruiz, N. Ungerleider, H. Nakhoul, E. F. Flemington, E. Kandil, S. B. Shah, S. D. Savkovic, Ulcerative colitis immune cell landscapes and differentially expressed gene signatures determine novel regulators and predict clinical response to biologic therapy. Sci. Rep. 11, 9010 (2021).

22. K. J. Eash, A. M. Greenbaum, P. K. Gopalan, D. C. Link, CXCR2 and CXCR4 antagonistically regulate neutrophil trafficking from murine bone marrow. J Clin Invest 120, 2423–2431 (2010).

23. N. Borregaard, K. Lollike, L. Kjeldsen, H. Sengeløv, L. Bastholm, M. H. Nielsen, D. F. Bainton, Human neutrophil granules and secretory vesicles. Eur. J. Haematol. 51, 187–198 (1993).

24. J. B. Cowland, N. Borregaard, The individual regulation of granule protein mRNA levels during neutrophil maturation explains the heterogeneity of neutrophil granules. J. Leukoc. Biol. 66, 989–995 (1999).

25. J. T. Chang, Pathophysiology of Inflammatory Bowel Diseases. N. Engl. J. Med. 383, 2652–2664 (2020).

26. T. Castro-Dopico, T. W. Dennison, J. R. Ferdinand, R. J. Mathews, A. Fleming, D. Clift, B. J. Stewart, C. Jing, K. Strongili, L. I. Labzin, E. J. M. Monk, K. Saeb-Parsy, C. E. Bryant, S. Clare, M. Parkes, M. R. Clatworthy. Anti-commensal IgG Drives Intestinal Inflammation and Type 17 Immunity in Ulcerative Colitis. Immunity 50, 1099–1114.e10 (2019).

27. M. Uzzan, J. C. Martin, L. Mesin, A. E. Livanos, T. Castro-Dopico, R. Huang, F. Petralia, G. Magri, S. Kumar, Q. Zhao, A. K. Rosenstein, M. Tokuyama, K. Sharma, R. Ungaro, R. Kosoy, D. Jha, J. Fischer, H. Singh, M. E. Keir, N. Ramamoorthi, W. E. O’Gorman, B. L. Cohen, A. Rahman, F. Cossarini, A. Seki, L. Leyre, S. T. Vaquero, S. Gurunathan, E. K. Grasset, B. Losic, M. Dubinsky, A. J. Greenstein, Z. Gottlieb, P. Legnani, J. George, H. Irizar, A. Stojmirovic, C. Brodmerkel, A. Kasarkis, B. E. Sands, G. Furtado, S. A. Lira, Z. K. Tuong, H. M. Ko, A. Cerutti, C. O. Elson, M. R. Clatworthy, M. Merad, M. Suárez-Fariñas, C. Argmann, J. A. Hackney, G. D. Victora, G. J. Randolph, E. Kenigsberg, J. F. Colombel, S. Mehandru. Ulcerative colitis is characterized by a plasmablast-skewed humoral response associated with disease activity. Nat. Med. 28, 766–779 (2022).

28. A. Kałużna, P. Olczyk, K. Komosińska-Vassev, The Role of Innate and Adaptive Immune Cells in the Pathogenesis and Development of the Inflammatory Response in Ulcerative Colitis. J. Clin. Med. 11, 400 (2022).

29. D. Muthas, A. Reznichenko, C. A. Balendran, G. Böttcher, I. G. Clausen, C. K. Mårdh, T. Ottosson, M. Uddin, T. T. MacDonald, S. Danese, M. B. Hansen, Neutrophils in ulcerative colitis: a review of selected biomarkers and their potential therapeutic implications. Scand. J. Gastroenterol. 52, 125–135 (2017).

30. M. D. Hu, K. L. Edelblum, Sentinels at the Frontline: the Role of Intraepithelial Lymphocytes in Inflammatory Bowel Disease. Curr. Pharmacol. Rep. 3, 321–334 (2017).

31. D. J. Gibson, E. J. Ryan, G. A. Doherty, Keeping the Bowel Regular. Inflamm. Bowel Dis. 19, 2716–2724 (2013).

32. R. Gaujoux, E. Starosvetsky, N. Maimon, F. Vallania, H. Bar-Yoseph, S. Pressman, R. Weisshof, I. Goren, K. Rabinowitz, M. Waterman, H. Yanai, I. Dotan, E. Sabo, Y. Chowers, P. Khatri, S. S. Shen-Orr, I. I. research N. (IIRN). Cell-centred meta-analysis reveals baseline predictors of anti-TNFα non-response in biopsy and blood of patients with IBD. Gut 68, 604 (2019).

33. B. Huang, Z. Chen, L. Geng, J. Wang, H. Liang, Y. Cao, H. Chen, W. Huang, M. Su, H. Wang, Y. Xu, Y. Liu, B. Lu, H. Xian, H. Li, H. Li, L. Ren, J. Xie, L. Ye, H. Wang, J. Zhao, P. Chen, L. Zhang, S. Zhao, T. Zhang, B. Xu, D. Che, W. Si, X. Gu, L. Zeng, Y. Wang, D. Li, Y. Zhan, D. Delfouneso, A. M. Lew, J. Cui, W. H. Tang, Y. Zhang, S. Gong, F. Bai, M. Yang, Y. Zhang, Mucosal Profiling of Pediatric-Onset Colitis and IBD Reveals Common Pathogenics and Therapeutic Pathways. Cell 179, 1160–1176.e24 (2019).

34. S. Vermeire, P. L. Lakatos, T. Ritter, S. Hanauer, B. Bressler, R. Khanna, K. Isaacs, S. Shah, A. Kadva, H. Tyrrell, Y. S. Oh, S. Tole, A. Chai, J. Pulley, C. Eden, W. Zhang, B. G. Feagan, L. S. Group, P. Abraham, M. A. C. Júnior, H. Aguilar, T. Ahmed, I. Altorjay, V. Andersen, R. Arai, H. Arnold, K. Ausk, J. Axler, K. Ayub, A. Balekuduru, G. B. Neto, I. Bassan, B. Behm, P. Bekal, S. Bhatia, B. Bod, C. E. B. Mello, J. Brandeburova, J. Breedt, B. Bressler, I. Chopey, M. Connor, R. Corlin, C. A. C. Hernandez, A. De, A. de S. Rolim, S. D. F. Boratto, T. Dixon, D. D. Poli, D. Dresner, G. A. D. Vall, M. Ebert, R. Ehehalt, A. Ertan, R. E. Valencia, J. Etzel, J. Fallingborg, B. Feagan, M. Fedurco, E. F. Castro, V. F. de A. Borges, M. Finklestein, A. Fischer, M. Fleisher, A. R. F. Rendon, R. Fogel, O. F. Junior, C. Freedland, D. Gatof, K. Gill, H. Glerup, V. Gokak, E. Goldin, H. A. G. Jaramillo, N. Gupta, Z. Gurzo, O. Gyrina, M. A. Habeeb, S. Hanauer, R. Hardi, W. Harlan, A. Hemaidan, M. Heyman, P. Hoffmann, W. Holderman, F. Holtkamp-Endemann, G. Horvat, K. Isaacs, E. Israeli, S. J. Miszputen, S. Jensen, K. Johnson, J. Jones, O. Junior, B. Kadleckova, M. Kalla, Z. Kallo, N. Karyotakis, L. Katz, L. Katz, N. Kaur, R. Khanna, P. Kohout, P. Lakatos, E. L. de los Reyes, R. Lee, B. Leman, O. Levchenko, H. Levine, L. L. B. C. Braga, E. Loftus, T. Lohdanidi, R. Longman, J. Lozano, C. Maaser, L. Madi-Szabo, E. F. Malluta, J. Marshall, F. M. Silva, K. Maynard, A. Meder, C. Mehta, P. Minarik, J. Mueller, S. Mukewar, B. Nagy, V. Neiko, M. Neurath, B. Nichol, J. Novick, N. Pai, W. Pandak, S. Panigrahi, U.-F. Pape, R. Paraná, N. Parekh, B. Patel, G. Pecsi, S. Peralta, M. Pesta, E. Peterfai, C. Petruzzellis, R. Petryka, R. Pica, C. Piniella, V. Pratha, V. Prochazka, S. Prokopchuk, L. Prystupa, A. Puri, T. Rainis, B. R. Kumar, O. R. Júnior, I. Rishko, T. Ritter, B. Robbins, E. Rock, M. R. Borba, M. Rodriguez, J. Rozciecha, A. Y. R. Flores, G. Rydzewska, R. Safadi, S. Saibeni, A. Schirbel, W. Schmiegel, R. Schnabel, H. Schneider, A. Segui, J. Seidelin, U. Seidler, J. Sellin, I. Shafran, S. Shah, A. Sheikh, A. Sherman, H. Shirin, A. Shukla, F. Siddiqui, R. Sike, A. Sood, A. Stallmach, M. Stanislavchuk, M. Staun, D. Stein, A. Steinberg, H. Steinhart, J. Stifft, R. Tandon, V. Tantry, S. Thiwan, M. Treiber, J. Ulbrych, J. Valentine, R. Vasudeva, B. Vaughn, B. Velasco, A. Vincze, M. Volfova, M. Waterman, L. M. Weiss, E. Wiesner, A. Williams, T. Witthoeft, R. Wohlman, J. Wright, J. K. Y. Furusho, Z. Younes, K. Yousuf, Y. Zborivskyy, S. Zeuzem, V. Zhdan, Etrolizumab for maintenance therapy in patients with moderately to severely active ulcerative colitis (LAUREL): a randomised, placebo-controlled, double-blind, phase 3 study. Lancet Gastroenterol. Hepatol. 7, 28–37 (2022).

35. L. Peyrin-Biroulet, A. Hart, P. Bossuyt, M. Long, M. Allez, P. Juillerat, A. Armuzzi, E. V. Loftus, E. Ostad-Saffari, A. Scalori, Y. S. Oh, S. Tole, A. Chai, J. Pulley, S. Lacey, W. J. Sandborn, H. S. Group, H. Aguilar, T. Ahmad, E. Akriviadis, X. A. Mante, M. Allez, I. Altorjay, A. Ananthakrishnan, V. Andersen, M. A. Garcia, A. Armuzzi, G. Aumais, I. Avni-Biron, J. Axler, K. Ayub, F. Baert, M. Bafutto, G. Bamias, I. Bassan, C. Baum, L. Beaugerie, B. Behm, P. Bekal, M. Bennett, F. B. S. Jose, C. Bernstein, D. Bettenworth, S. Bhaskar, L. Biancone, B. Bilir, M. Blaeker, S. Bloom, V. Bohman, F. J. B. Padilla, P. Bossuyt, Y. Bouhnik, G. Bouma, R. Bourdages, S. Brand, B. Bressler, M. Brückner, C. Buening, F. Carbonnel, T. Caves, J. Chapman, J. H. Cheon, N. Chiba, C. Chioncel, D. Christodoulou, M. Clodi, A. Cohen, G. R. Corazza, R. Corlin, R. Cosintino, F. Cummings, R. Dalal, S. Danese, M. D. Maeyer, C. F. D. M. Francesconi, A. D. Silva, H. Debinski, P. Desreumaux, O. Dewit, G. D’Haens, S. D. F. Boratto, J. N. Ding, T. Dixon, G. Dryden, G. A. D. Vall, M. Ebert, A. E. Piudo, R. Ehehalt, M. Elkhashab, C. Ennis, J. Etzel, J. Fallingborg, B. Feagan, R. Fejes, D. F. de C. Mazo, V. F. de A. Borges, A. Fischer, A. Fixelle, M. Fleisher, S. Fowler, B. Freilich, K. Friedenberg, W. Fries, C. Fulop, M. Fumery, S. Fuster, G. G. Kiss, S. G. Lopez, S. Gassner, K. Gill, C. G. de S. Joseph, P. Ginsburg, P. Gionchetti, E. Goldin, A.-E. Goldis, H. A. G. Jaramillo, M. Gonciarz, G. Gordon, D. Green, J.-C. Grimaud, R. G. Rodriguez, Z. Gurzo, A. Gutierrez, T. Gyökeres, K. B. Hahm, S. Hanauer, J. Hanson, W. H. III, P. Hasselblatt, B. Hayee, X. Hebuterne, P. Hendy, M. Heyman, P. Higgins, R. Hilal, P. Hindryckx, F. Hoentjen, P. Hoffmann, F. Holtkamp-Endemann, G. Holtmann, G. Horvat, S. Howaldt, S. Huber, I. Ibegbu, M. I. I. Colomino, P. Irving, K. Isaacs, K. Jagarlamudi, R. Jain, S. J. Miszputen, J. Jansen, J. Jones, P. Juillerat, J. Karagiannis, N. Karyotakis, A. Kaser, L. Katz, S. Katz, L. Katz, N. Kaur, E. Kazenaite, R. Khanna, S. Khurana, J. S. Kim, Y.-H. Kim, S. K. Kim, D. Kim, J. Klaus, D. Kleczkowski, P. Kohout, B. Korczowski, G. Kouklakis, I. Koutroubakis, R. Krause, T. Kristof, I. Kronborg, A. Krummenerl, L. Kupcinskas, J. L. Molteni, D. Laharie, A. Lahat-zok, J. Lee, K.-M. Lee, R. Leong, H. Levine, J. Limdi, J. Lindsay, N. Lodhia, E. Loftus, R. Longman, P. L. Serrano, E. Louis, M. H. L. Pereira, J. Lowe, S. Lueth, M. Lukas, G. Maconi, F. Macrae, L. Madi-Szabo, U. Mahadevan-Velayos, E. F. Malluta, F. Mana, P. Mannon, G. Mantzaris, I. M. Jimenez, M. D. M. Arranz, R.-B. Mateescu, F. Mazzoleni, A. Meder, E. Melzer, J. Mertens, K. Mimidis, B. Mitchell, T. Molnar, G. Moore, L. A. M. Garza, R. Mountifield, V. Muls, C. Murray, B. Nagy, M. Neurath, A. Nguyen, R. Panaccione, W. Pandak, J. P. Diaz, J. Park, L. Pastorelli, B. Patel, M. Peck-Radosavljevic, G. Pecsi, F. Peerani, J. P. Gisbert, M. Pesta, R. Petryka, L. Peyrin-Biroulet, R. Phillips, M. Pierik, V. Pratha, V. Prochazka, I. Racz, G. Radford-Smith, D. R. Castañeda, O. R. Júnior, J. Regula, J.-M. Reimund, B. Robbins, X. Roblin, F. Rogai, G. Rogler, J. Rozciecha, D. Rubin, A. Y. R. Flores, M. Rupinski, G. Rydzewska, S. Saha, S. Saibeni, A. Salamon, Z. Sallo, B. Salzberg, D. Samuel, S. Samuel, W. Sandborn, E. V. Savarino, A. Schirbel, R. Schnabel, S. Schreiber, J. Scott, S. Sedghi, F. Seibold, J. Seidelin, U. Seidler, A. Shaban, I. Shafran, A. Sheikh, A. Sherman, H. Shirin, P. Smolinski, G. A. Song, K. Soufleris, A. Speight, D. Staessen, A. Stallmach, M. Staun, D. Stein, H. Steinhart, J. Stifft, D. Stokesberry, A. Sturm, K. Sultan, G. Szekely, K. Tagore, H. Tanno, L. Thin, S. Thiwan, C. Thomas, M. Tichy, G. T. Toth, Z. Tulassay, J. Ulbrych, J. Valentine, M. Varga, E. Vasconcellos, A. Vaughn, B. Velasco, F. Velazquez, S. Vermeire, E. Villa, A. Vincze, H. Vogelsang, M. Volfova, L. Vuitton, P. Vyhnalek, P. Wahab, J. Walldorf, M. Waterman, J. Weber, L. M. Weiss, A. Wiechowska-Kozlowska, E. Wiesner, T. Witthoeft, R. Wohlman, B. Wozniak-Stolarska, B. Yacyshyn, B.-D. Ye, Z. Younes, L. Y. Sassaki, C. Zaltman, S. Zeuzem, Etrolizumab as induction and maintenance therapy for ulcerative colitis in patients previously treated with tumour necrosis factor inhibitors (HICKORY): a phase 3, randomised, controlled trial. Lancet Gastroenterol. Hepatol. 7, 128–140 (2022).

36. S. Danese, J.-F. Colombel, M. Lukas, J. P. Gisbert, G. D’Haens, B. Hayee, R. Panaccione, H.-S. Kim, W. Reinisch, H. Tyrrell, Y. S. Oh, S. Tole, A. Chai, K. Chamberlain-James, M. T. Tang, S. Schreiber, G. S. Group, N. Aboo, T. Ahmad, X. A. Mante, M. Allez, S. Almer, R. Altwegg, M. A. Garcia, R. Arasaradnam, S. Ardizzone, A. Armuzzi, I. Arnott, G. Aumais, I. Avni-Biron, P. Barrow, I. Beales, F. B. S. Jose, A. Bezuidenhout, L. Biancone, M. Blaeker, S. Bloom, B. Bokemeyer, F. Bossa, P. Bossuyt, G. Bouguen, Y. Bouhnik, G. Bouma, R. Bourdages, A. Bourreille, C. Boustiere, T. Brabec, S. Brand, C. Buening, A. Buisson, G. Cadiot, X. C. Calvo, F. Carbonnel, D. Carpio, J. H. Cheon, N. Chiba, C. Chioncel, N.-C. Cimpoeru, M. Clodi, G. R. Corazza, R. Cosintino, J. Cotter, T. Creed, F. Cummings, S. Danese, G. L. de’ Angelis, M. D. Maeyer, M. Desai, E. Desilets, P. Desreumaux, O. Dewit, G. D’Haens, J. Dinter, E. D. Dobru, T. Douda, D. L. Dumitrascu, M. Ebert, A. E. Piudo, M. Elkhashab, C. S. Eun, B. Feagan, R. Fejes, A. Fidalgo, S. Fishman, B. Flourié, S. Fowler, W. Fries, C. Fulop, M. Fumery, G. G. Kiss, S. Gassner, D. Gaya, B. Germanà, L. S. Gheorghe, C. G. de S. Joseph, P. Gionchetti, A.-E. Goldis, R. Gonçalves, J.-C. Grimaud, T. Gyökeres, H. Hagege, A. Haidar, H. Hartmann, P. Hasselblatt, B. Hayee, X. Hebuterne, P. Hellström, P. Hindryckx, H. Hlavova, F. Hoentjen, S. Howaldt, L. Hrdlicka, K. C. Huh, M. I. I. Colomino, F. Ionita-Radu, P. Irving, J. Jahnsen, B. Jang, J. Jansen, S. W. Jeon, R. J. Martinez, P. Juillerat, P. Karlén, A. Kaser, R. Keil, D. Kejariwal, D. Keret, R. Khanna, D. Kim, D. H. Kim, H.-J. Kim, H.-S. Kim, J. S. Kim, K. Kim, K.-J. Kim, S. K. Kim, Y.-H. Kim, J. Klaus, A. Kohn, V. Kojecky, J. S. Koo, R. Kozak, M. Kremer, T. Kristof, F. Kruger, D. Laharie, A. Lahat-zok, E. Landa, J. Lee, K.-M. Lee, K. L. Lee, Y. Lee, F. Lenze, W. C. Lim, J. Limdi, J. Lindsay, P. L. Serrano, E. Louis, S. Lueth, M. Lukas, G. Maconi, F. Mana, S. Mann, J. Mansfield, S. Marchi, M. Marino, J. Marshall, M. D. M. Arranz, R.-B. Mateescu, J. McLaughlin, S. McLaughlin, E. Melzer, J. Mertens, P. Mitrut, T. Molnar, V. Muls, P. Munuswamy, C. Murray, T. Naftali, V. Naidoo, Y. Nanabhay, L. Negreanu, A. Nguyen, T. Ochsenkuehn, A. Orlando, R. Panaccione, J. P. Diaz, M. Paritsky, D. I. Park, J. Park, L. Pastorelli, M. Peck-Radosavljevic, F. Peerani, J. P. Gisbert, L. Peyrin-Biroulet, L. Picon, M. Pierik, T. Ponich, F. Portela, M. J. Prins, I. Racz, K. F. Rahman, J.-M. Reimund, M. Reinshagen, X. Roblin, R. Rocca, F. Rogai, G. Rogler, A. Salamon, E. Salazar, Z. Sallo, S. Samuel, M. de los S. S. Cuffi, E. V. Savarino, V. Savarino, G. Savoye, S. Schreiber, A. Seicean, C. Selinger, D. M. Serra, H. H. Shim, S. Shin, B. Siegmund, J. Siffledeen, W. Simmonds, J. Smid, J. Sollano, G. A. Song, A. Speight, I. Sporea, A. Staessen, G. Stancu, A. Steel, D. Stepek, V. Stoica, A. Sturm, G. Szekely, T. K. Tan, C. T. Samso, J. Thomson, M. Tichy, G. T. Toth, Z. Tulassay, M. Vangeli, M. Varga, A. Vieira, S. Viennot, E. Villa, P. Vitek, H. Vogelsang, P. Vyhnalek, P. Wahab, J. Walldorf, B. D. Ye, C. Ziady, Etrolizumab versus infliximab for the treatment of moderately to severely active ulcerative colitis (GARDENIA): a randomised, double-blind, double-dummy, phase 3 study. Lancet Gastroenterol. Hepatol. 7, 118–127 (2022).

37. N. R. West, A. N. Hegazy, B. M. J. Owens, S. J. Bullers, B. Linggi, S. Buonocore, M. Coccia, D. Görtz, S. This, K. Stockenhuber, J. Pott, M. Friedrich, G. Ryzhakov, F. Baribaud, C. Brodmerkel, C. Cieluch, N. Rahman, G. Müller-Newen, R. J. Owens, A. A. Kühl, K. J. Maloy, S. E. Plevy, S. Keshav, S. P. L. Travis, F. Powrie, C. Arancibia, A. Bailey, E. Barnes, B. Bird-Lieberman, O. Brain, B. Braden, J. Collier, J. East, L. Howarth, S. Keshav, P. Klenerman, S. Leedham, R. Palmer, F. Powrie, A. Rodrigues, A. Simmons, P. Sullivan, H. Uhlig, Oncostatin M drives intestinal inflammation and predicts response to tumor necrosis factor–neutralizing therapy in patients with inflammatory bowel disease. Nat Med 23, 579–589 (2017).

38. C. Zhang, W. Shu, G. Zhou, J. Lin, F. Chu, H. Wu, Z. Liu. Anti-TNF-α Therapy Suppresses Proinflammatory Activities of Mucosal Neutrophils in Inflammatory Bowel Disease. Mediators Inflamm. 2018 Nov 22;2018:3021863.

39. G. W. Tew, J. A. Hackney, D. Gibbons, C. A. Lamb, D. Luca, J. G. Egen, L. Diehl, J. E. Anderson, S. Vermeire, J. C. Mansfield, B. G. Feagan, J. Panes, D. C. Baumgart, S. Schreiber, I. Dotan, W. J. Sandborn, J. A. Kirby, P. M. Irving, G. D. Hertogh, G. A. V. Assche, P. Rutgeerts, S. O’Byrne, A. Hayday, M. E. Keir, S. O’Byrne, A. Hayday, M. E. Keir, Association Between Response to Etrolizumab and Expression of Integrin αE and Granzyme A in Colon Biopsies of Patients With Ulcerative Colitis. Gastroenterology 150, 477–487.e9 (2016).

40. R. Ungaro, J.-F. Colombel, T. Lissoos, L. Peyrin-Biroulet, A Treat-to-Target Update in Ulcerative Colitis: A Systematic Review. Am. J. Gastroenterol. 114, 874–883 (2019).

41. S. Danese, G. Roda, L. Peyrin-Biroulet, Evolving therapeutic goals in ulcerative colitis: towards disease clearance. Nat. Rev. Gastroenterol. Hepatol. 17, 1–2 (2020).

42. R. Grieshaber-Bouyer, F. A. Radtke, P. Cunin, G. Stifano, A. Levescot, B. Vijaykumar, N. Nelson-Maney, R. B. Blaustein, P. A. Monach, P. A. Nigrovic, O. Aguilar, R. Allan, J. Astarita, K. F. Austen, N. Barrett, A. Baysoy, C. Benoist, B. D. Brown, M. Buechler, J. Buenrostro, M. A. Casanova, K. Chowdhary, M. Colonna, T. Crowl, T. Deng, F. Desland, M. Dhainaut, J. Ding, C. Dominguez, D. Dwyer, M. Frascoli, S. Gal-Oz, A. Goldrath, T. Johanson, S. Jordan, J. Kang, V. Kapoor, E. Kenigsberg, J. Kim, K. wook Kim, E. Kiner, M. Kronenberg, L. Lanier, C. Laplace, C. Lareau, A. Leader, J. Lee, A. Magen, B. Maier, A. Maslova, D. Mathis, A. McFarland, M. Merad, A. Meunier, P. A. Monach, S. Mostafavi, S. Muller, C. Muus, H. Ner-Gaon, Q. Nguyen, G. Novakovsky, S. Nutt, K. Omilusik, A. Ortiz-Lopez, M. Paynich, V. Peng, M. Potempa, R. Pradhan, S. Quon, R. Ramirez, D. Ramanan, G. Randolph, A. Regev, S. A. Rose, K. Seddu, T. Shay, A. Shemesh, J. Shyer, C. Smilie, N. Spidale, A. Subramanian, K. Sylvia, J. Tellier, S. Turley, B. Vijaykumar, A. Wagers, C. Wang, P. L. Wang, A. Wroblewska, L. Yang, A. Yim, H. Yoshida, The neutrotime transcriptional signature defines a single continuum of neutrophils across biological compartments. Nature Communications 12, 1–21 (2021).

43. J. M. Adrover, C. del Fresno, G. Crainiciuc, M. I. Cuartero, M. Casanova-Acebes, L. A. Weiss, H. Huerga-Encabo, C. Silvestre-Roig, J. Rossaint, I. Cossío, A. V. Lechuga-Vieco, J. García-Prieto, M. Gómez-Parrizas, J. A. Quintana, I. Ballesteros, S. Martin-Salamanca, A. Aroca-Crevillen, S. Z. Chong, M. Evrard, K. Balabanian, J. López, K. Bidzhekov, F. Bachelerie, A. Abad-Santos, C. Muñoz-Calleja, A. Zarbock, O. Soehnlein, C. Weber, L. G. Ng, C. Lopez-Rodriguez, D. Sancho, M. A. Moro, B. Ibáñez, A. Hidalgo, A Neutrophil Timer Coordinates Immune Defense and Vascular Protection. Immunity 50, 390–402.e10 (2019).

44. I. Dotan, L. Werner, S. Vigodman, S. Weiss, E. Brazowski, N. Maharshak, O. Chen, H. Tulchinsky, Z. Halpern, H. Guzner-Gur, CXCL12 Is a constitutive and inflammatory chemokine in the intestinal immune system. Inflamm. Bowel Dis. 16, 583–592 (2010).

45. T. H. Schmidt, O. Bannard, E. E. Gray, J. G. Cyster, CXCR4 promotes B cell egress from Peyer’s patches. J. Exp. Med. 210, 1099–1107 (2013).

46. L. Werner, H. Elad, E. Brazowski, H. Tulchinsky, S. Vigodman, U. Kopylov, Z. Halpern, H. Guzner-Gur, I. Dotan, Reciprocal regulation of CXCR4 and CXCR7 in intestinal mucosal homeostasis and inflammatory bowel disease. J. Leukoc. Biol. 90, 583–590 (2011).

47. P. Czarnewski, S. M. Parigi, C. Sorini, O. E. Diaz, S. Das, N. Gagliani, E. J. Villablanca, Conserved transcriptomic profile between mouse and human colitis allows unsupervised patient stratification. Nat. Commun. 10, 2892 (2019).

48. F. Zhu, H. He, L. Fan, C. Ma, Z. Xu, Y. Xue, Y. Wang, C. Zhang, G. Zhou, Blockade of CXCR2 suppresses proinflammatory activities of neutrophils in ulcerative colitis. Am. J. Transl. Res. 12, 5237–5251 (2020).

49. S. M. Farooq, R. Stillie, M. Svensson, C. Svanborg, R. M. Strieter, A. W. Stadnyk, Therapeutic Effect of Blocking CXCR2 on Neutrophil Recruitment and Dextran Sodium Sulfate-Induced Colitis. J. Pharmacol. Exp. Ther. 329, 123–129 (2009).

50. A. F. Bento, D. F. P. Leite, R. F. Claudino, D. B. Hara, P. C. Leal, J. B. Calixto, The selective nonpeptide CXCR2 antagonist SB225002 ameliorates acute experimental colitis in mice. J. Leukoc. Biol. 84, 1213–1221 (2008).

51. S. Sitaru, A. Budke, R. Bertini, M. Sperandio, Therapeutic inhibition of CXCR1/2: where do we stand? Intern. Emerg. Med. 18, 1647–1664 (2023).

52. L. Mao, A. Kitani, M. Similuk, A. J. Oler, L. Albenberg, J. Kelsen, A. Aktay, M. Quezado, M. Yao, K. Montgomery-Recht, I. J. Fuss, W. Strober, Loss-of-function CARD8 mutation causes NLRP3 inflammasome activation and Crohn’s disease. J. Clin. Investig. 128, 1793–1806 (2018).

53. E. Shaul, M. A. Conrad, N. Dawany, T. Patel, M. C. Canavan, A. Baccarella, S. Weinbrom, D. Aleynick, K. E. Sullivan, J. R. Kelsen, Canakinumab for the treatment of autoinflammatory very early onset- inflammatory bowel disease. Front. Immunol. 13, 972114 (2022).

54. G. Pau, J. Reeder, HTSeqGenie: A NGS analysis pipeline (2023; https://bioconductor.org/packages/HTSeqGenie).

55. T. D. Wu, S. Nacu, Fast and SNP-tolerant detection of complex variants and splicing in short reads. Bioinformatics 26, 873–881 (2010).

56. Y. Hao, S. Hao, E. Andersen-Nissen, W. M. Mauck, S. Zheng, A. Butler, M. J. Lee, A. J. Wilk, C. Darby, M. Zager, P. Hoffman, M. Stoeckius, E. Papalexi, E. P. Mimitou, J. Jain, A. Srivastava, T. Stuart, L. M. Fleming, B. Yeung, A. J. Rogers, J. M. McElrath, C. A. Blish, R. Gottardo, P. Smibert, R. Satija, Integrated analysis of multimodal single-cell data. Cell 184, 3573–3587.e29 (2021).

57. I. Korsunsky, N. Millard, J. Fan, K. Slowikowski, F. Zhang, K. Wei, Y. Baglaenko, M. Brenner, P. Loh, S. Raychaudhuri, Fast, sensitive and accurate integration of single-cell data with Harmony. Nat. Methods 16, 1289–1296 (2019).

58. G. Heimberg, T. Kuo, D. DePianto, T. Heigl, N. Diamant, O. Salem, G. Scalia, T. Biancalani, S. Turley, J. Rock, H. C. Bravo, J. Kaminker, J. A. V. Heiden, A. Regev, Scalable querying of human cell atlases via a foundational model reveals commonalities across fibrosis-associated macrophages. bioRxiv 2023.07.18.549537 (2023).

59. L. Kong, V. Pokatayev, A. Lefkovith, G. T. Carter, E. A. Creasey, C. Krishna, S. Subramanian, B. Kochar, O. Ashenberg, H. Lau, A. N. Ananthakrishnan, D. B. Graham, J. Deguine, R. J. Xavier, The landscape of immune dysregulation in Crohn’s disease revealed through single-cell transcriptomic profiling in the ileum and colon. Immunity 56, 444–458.e5 (2023).

60. M. D. Robinson, D. J. McCarthy, G. K. Smyth, edgeR: a Bioconductor package for differential expression analysis of digital gene expression data. Bioinformatics 26, 139–140 (2010).

61. I. Korsunsky, A. Nathan, N. Millard, S. Raychaudhuri, Presto scales Wilcoxon and auROC analyses to millions of observations. bioRxiv, 653253 (2019).

62. C. Trapnell, D. Cacchiarelli, J. Grimsby, P. Pokharel, S. Li, M. Morse, N. J. Lennon, K. J. Livak, T. S. Mikkelsen, J. L. Rinn, The dynamics and regulators of cell fate decisions are revealed by pseudotemporal ordering of single cells. Nat. Biotechnol. 32, 381–386 (2014).

63. X. Qiu, Q. Mao, Y. Tang, L. Wang, R. Chawla, H. A. Pliner, C. Trapnell, Reversed graph embedding resolves complex single-cell trajectories. Nat. Methods 14, 979–982 (2017).

64. J. Cao, M. Spielmann, X. Qiu, X. Huang, D. M. Ibrahim, A. J. Hill, F. Zhang, S. Mundlos, L. Christiansen, F. J. Steemers, C. Trapnell, J. Shendure, The single-cell transcriptional landscape of mammalian organogenesis. Nature 566, 496–502 (2019).

65. S. Jin, C. F. Guerrero-Juarez, L. Zhang, I. Chang, R. Ramos, C.-H. Kuan, P. Myung, M. V. Plikus, Q. Nie, Inference and analysis of cell-cell communication using CellChat. Nat. Commun. 12, 1088 (2021).

66. R. Casadio, E. Frigimelica, P. Bossù, D. Neumann, M. U. Martin, A. Tagliabue, D. Boraschi, Model of interaction of the IL-1 receptor accessory protein IL-1RAcP with the IL-1β/IL-1RI complex. FEBS Lett. 499, 65–68 (2001).

